# A comprehensive benchmark and guide for sequence-function interpretable deep learning models in genomics

**DOI:** 10.1101/2025.01.06.631405

**Authors:** Canzhuang Sun, Yu Sun, Kang Xu, Zhijie He, Hao Li, Yaru Li, Zongyuan Yu, Yuyang Wang, Xuanwei Lin, Xiang Xu, Pengzhen Hu, Xiaochen Bo, Mingzhi Liao, Hebing Chen

**Affiliations:** College of Life Sciences, Center of Bioinformatics, Northwest A&F University, China; Academy of Military Medical Sciences; College of Computer and Data Science, Fuzhou University, China; School of Software, Shandong University, China

## Abstract

The development of sequence-based deep learning methods has greatly increased our understanding of how sequence determines function. In parallel, numerous interpretable algorithms have been developed to address complex tasks, such as elucidating sequence regulatory syntax and analyzing non-coding variants from trained models. However, few studies have systematically compared and evaluated the performance and interpretability of these algorithms. Here, we introduce a comprehensive benchmark framework for evaluating sequence-to-function models. We systematically evaluated multiple models and DNA language foundation models using 369 ATAC-seq datasets, employing diverse training strategies and evaluation metrics to uncover their critical strengths and limitations. Our benchmark study highlights that different model architectures and interpretability methods are better suited to specific scenarios. Negative samples derived from naturally inactive regions outperform synthetic sequences, whereas single-cell tasks require specialized models. Additionally, we demonstrate that interpretable sequence-function models can complement traditional sequence alignment methods in studying cross-species enhancer regulatory logic. We also provide a pipeline to help researchers select the optimal sequence-function prediction and interpretability algorithms.

## Introduction

Cis-regulatory elements are crucial for the dynamic regulation of the genome in development, cellular homeostasis, external stimuli responses, and disease^1^. These elements overlap significantly with trait-associated genetic variants, and their disruption is linked to genetic disorders and cancer. Chromatin accessibility often marks putative cis-regulatory elements, such as promoters and enhancers, providing a map of the regulatory landscape^2, 3^. However, interpreting the sequence determinants of this landscape remains challenging.

Recent advances in artificial intelligence and large-scale functional genomics have facilitated the development of sequence-to-function models, offering powerful tools to uncover the complex relationships between genomic sequences and regulatory functions in disease-relevant contexts^4–7^. These models, such as Basset, DanQ, DeepSEA, and Enformer, predict chromatin accessibility, transcription factor binding, histone modifications, and 3D genome folding, among other regulatory features^4, 6–11^. By employing interpretability techniques, these models can uncover cell type-specific regulatory grammars, enabling predictions of how sequence motifs influence regulatory landscapes^8, 12–16^. These models, trained on specific datasets, facilitate the design of efficient regulatory sequences, such as promoters and enhancers^17–19^. Moreover, they provide a valuable metric for sequencing data quality and enhance analyses of single-cell ATAC-seq data, supporting applications like cell clustering and batch effect correction^20, 21^.

Most sequence-to-function methods rely on convolutional neural networks (CNNs) or their variants, where convolutional filters scan sequences in a manner similar to position weight matrices (PWMs), and deeper layers aggregate information across larger regions^22^. Recently, long-sequence-based models using self-attention mechanisms have gained attention for their ability to capture long-range interactions, enabling genome-wide predictions of chromatin accessibility and 3D genome folding^23–26^. However, these complex models present challenges in extracting biologically meaningful insights and require further development of interpretability techniques to enhance their practical utility^27, 28^. In contrast, single-peak approaches are computationally efficient, well-suited for learning sequence regulatory syntax, and leverage existing interpretability methods to provide intuitive insights into evolutionary and developmental processes^29^. Despite their advantages, single-peak methods exhibit variability in design parameters, such as model architecture, sequence length, and sample selection, and lack a unified framework for evaluation. In the context of single-cell ATAC-seq data analysis, the applicability of models beyond the scBasset architecture remains uncertain^21^.

To address challenges in the predictive performance and interpretability of sequence-function models, we propose a unified framework for systematically evaluating existing deep learning approaches. This framework enables comprehensive comparisons of predictive accuracy and interpretability, with a particular focus on chromatin accessibility prediction. This framework enables comprehensive comparisons of predictive accuracy and interpretability, focusing on chromatin accessibility prediction. By identifying key strengths in model architectures, training strategies, and data processing techniques, we highlight critical factors affecting model performance. We developed a novel interpretability method to quantify key sequence regions, offering new insights into non-coding variants and transcription factor cooperativity. Our analysis also reveals regulatory element conservation during early vertebrate development, demonstrating the potential of interpretable models to complement sequence alignment methods in elucidating cross-species enhancer logic.

## Results

### Benchmark pipeline

In this benchmark study, we evaluated the performance of 11 algorithms in predicting chromatin accessibility. Among these, six algorithms—DeepSEA, Basset, DanQ, ExplaiNN, SATORI, and Scover—were specifically designed for diverse predictive tasks. Three models were DNA foundation models, namely DNABERT2^30, 31^, Nucleotide Transformer (NT) ^32^, and HyenaDNA^33^, while the remaining two were Transformer-based hybrid models (CNN+Transformer and CNN+Attention) (**Additional file FigS1**). To comprehensively assess their performance, we collected 369 ATAC-seq datasets from the ENCODE database, covering a broad range of tissues and cell lines. We examined model performance across various training strategies, including single-task versus multi-task learning, and regression versus binary classification tasks. Evaluations were conducted across multiple application scenarios, such as chromatin activity prediction and model interpretability. Following widely accepted data partitioning strategies, we designated chromosomes 8 and 9 as the test set, while the remaining chromosomes were used for training. For binary classification tasks, we employed AUROC and AUPRC as evaluation metrics, whereas Pearson correlation coefficient (PCC) and Mean Squared Error (MSE) were used for regression tasks. We evaluated the two learning strategies on single-task datasets and similarly assessed them on multi-task datasets. (Fig1.a).

**Fig 1.**
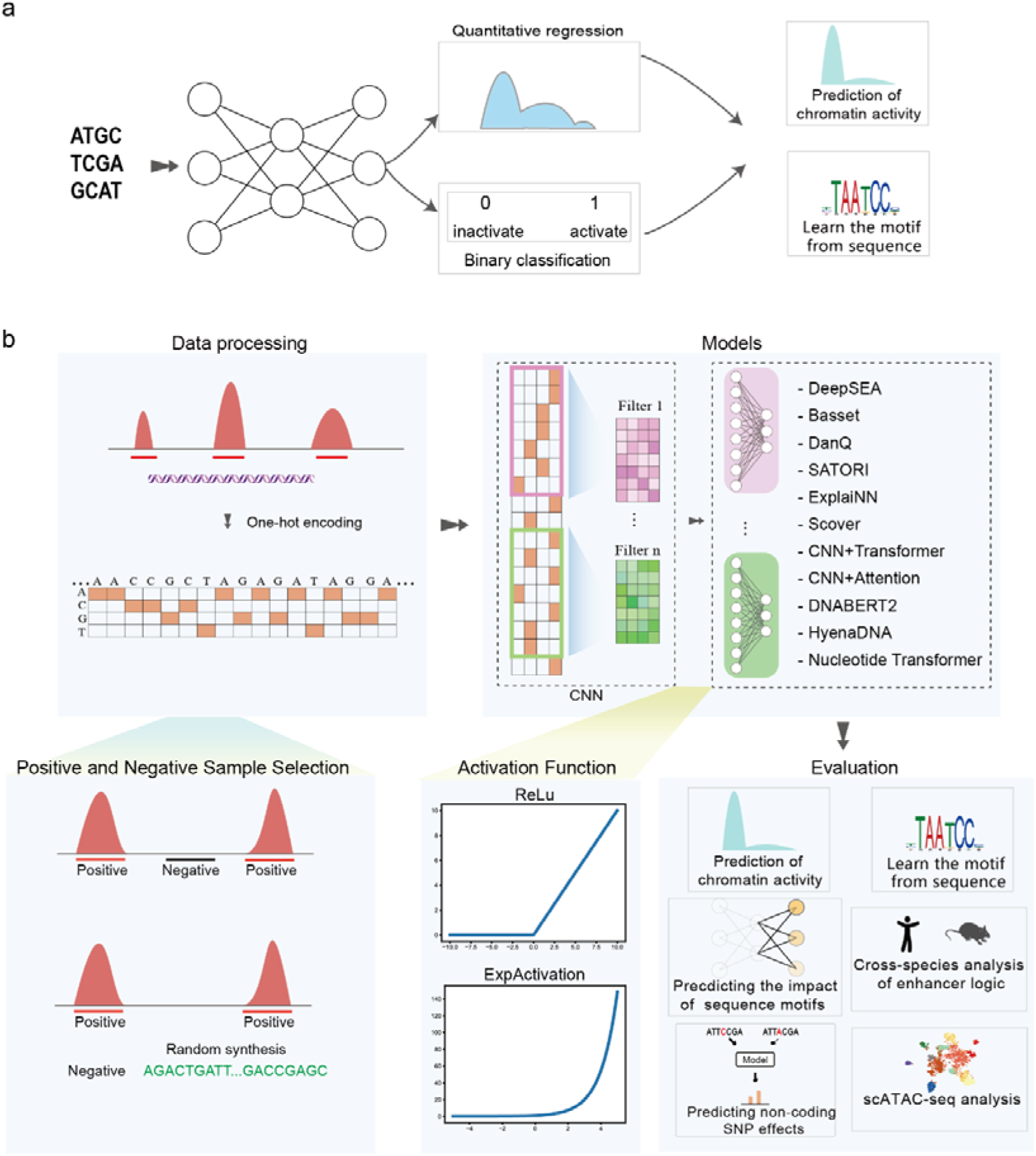
Overview of benchmarking framework. **a,** Comparison of binary and quantitative prediction tasks in regulatory genomics. **b,** Illustration of the three main components of sequence-function models: data preprocessing (input size, dataset construction), model training (activation functions), and evaluation (accuracy, robustness, interpretability, and biological scenario application)

To better understand the factors influencing the performance of chromatin accessibility prediction models and to utility in functional genomics of interpretable methods in downstream analyses, we developed a Python-based software package, cisFinder (**c**omprehensive **i**nterpretable **s**equence-**F**unct**i**o**n** mo**d**el **e**valuato**r**) (Fig1.b). Implemented using the Pytorch framework, **cisFinder** integrates most of the currently published deep learning methods for predicting epigenomic signals from sequence data. This tool supports a wide range of functions, including data preprocessing, model training, model interpretation, and performance evaluation. Using **cisFinder**, we further examined the effects of key parameters, such as input sequence length, positive and negative sample construction methods, activation functions, and model interpretability. Finally, we extended the application of these methods to specific research scenarios, demonstrating their versatility and potential in functional genomics. By providing researchers with a robust evaluation framework, **cisFinder** enables informed decisions about the most appropriate methods for diverse genomic studies.

### Performance of prediction algorithm for chromatin accessibility

Sequence-based approaches in functional genomics rely on model performance on independent test datasets to ensure the learned mapping reflects genuine sequence-function relationships rather than overfitting to the training data. To ensure a fair comparison, we used cisFinder to train all models with identical input sequence lengths and hyperparameters (input sequence length: 600 bp, learning rate: 0.001, batch size: 128).

In the single-task learning setting, most methods performed well, achieving AUROC and AUPRC scores exceeding 0.90 (Fig2.a). Among these, the LSTM-based models, DanQ and SATORI, exhibited exceptional performance, achieving mean AUROC/AUPRC scores of 0.964/0.961 and 0.960/0.956, respectively. DeepSEA also demonstrated strong performance, with AUROC/AUPRC values of 0.946/0.949. In comparison, Basset showed slightly lower performance, with AUROC/AUPRC scores of 0.925/0.926. Both models are based on fully convolutional architectures; however, DeepSEA contains a larger number of parameters than Basset. The Transformer-based CNN+Transformer outperformed the CNN+Attention model that utilizes attention mechanisms. ExplaiNN achieved robust performance, with AUROC and AUPRC values of 0.932 and 0.928, respectively. The simplified Scover architecture, comprising a single convolutional and linear layer, achieved the lowest performance (AUROC/AUPRC = 0.921/0.917). In the multi-task learning setting, DanQ and SATORI remained the top performers, while the relative performance trends of other models were consistent with those observed in the single-task scenario (Fig2.b). We assessed the convergence speed and overfitting tendencies of each model. With a fixed batch size of 128, DanQ, DeepSEA, Basset, ExplaiNN, and CNN+Transformer demonstrated rapid convergence within 10 epochs (FigS2.a). Although SATORI exhibited superior performance, its convergence speed was slower. All models showed varying degrees of overfitting, with the LSTM-based architectures, DanQ and SATORI, displaying relatively lower levels of overfitting (FigS2.b, FigS2.c). Consistent with previous studies, we observed a decline in the performance of deep learning models in cell type-specific regions (FigS2.d)^34^, with models performing better in regions associated with active histone modifications, such as H3K4me3 and H3K27ac, compared to repressive modifications like H3K9me3 and H3K27me3 (FigS2.e).

**Fig 2.**
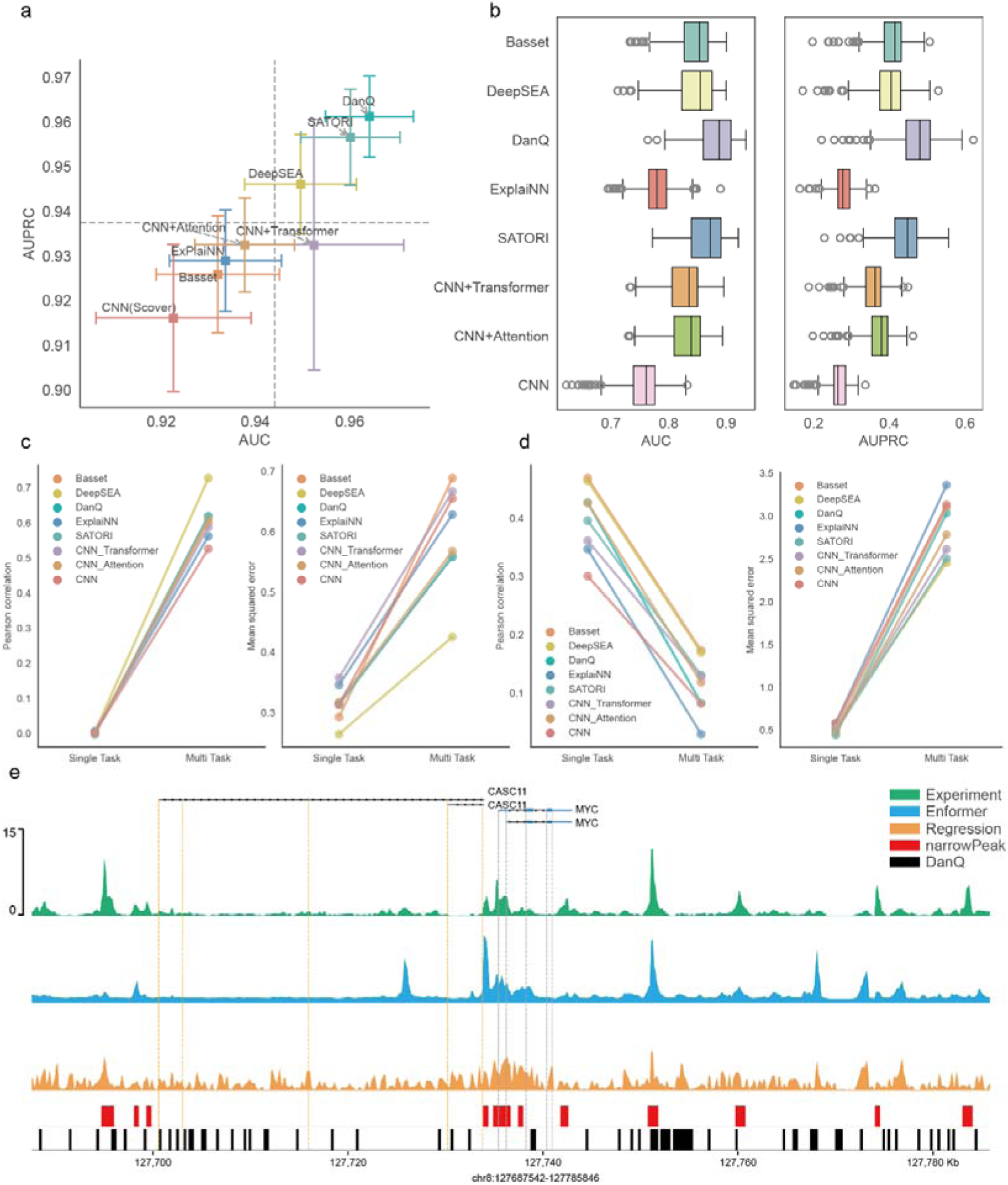
Performance of the algorithm in predicting binary chromatin accessibility. **a,** AUROC and AUPRC scores of single-task learning models on chromatin accessibility prediction. **b,** AUROC and AUPRC scores of multi-task learning models. **c,** Comparison of single-task regression and multi-task regression learning models on single-task datasets. **d,** Comparison of single-task regression and multi-task regression learning models on multi-task datasets. **e,** Performance of long-sequence modeling approaches (Enformer) compared with single-peak regression and classification methods.

We also compared the performance of DNA foundation models based on large language models (LLMs), including DNABERT2, Nucleotide Transformer (NT), HyenaDNA, and DanQ on ATAC-seq datasets from six cell lines. DanQ significantly outperformed the other three methods, indicating that purpose-built models are more effective than DNA language models in predicting chromatin accessibility (FigS2.f). One potential strategy for improving LLMs would be to fine-tune them on large-scale chromatin accessibility datasets. Additionally, we investigated the impact of single-task versus multi-task learning on chromatin accessibility prediction. When multi-task models were applied to single-task datasets, their performance was lower than that of single-task models. Notably, single-task models also outperformed multi-task models when evaluated on multi-task datasets (FigS3.a, Fig. S3.b). This discrepancy may arise from the ability of single-task models to better capture sequence features specific to individual cell types or tissues, whereas the trade-off in multi-task learning could reduce performance across tasks.

We evaluated the performance of various models on chromatin accessibility regression tasks using 369 ATAC-seq signal values (read counts) from the ENCODE database. Similar to classification tasks, we compared the performance of single-task and multi-task models. In single-task learning, Basset and DeepSEA achieved the highest performance, with average Pearson correlation coefficients (PCCs) of 0.43 and 0.42, respectively. DanQ followed closely (PCC 0.43) and demonstrated a higher potential performance ceiling. CNN+Attention performed slightly worse than DanQ (PCC 0.42), while all other models exhibited PCCs below 0.4 (FigS4.a). Although DanQ’s PCC was slightly lower than those of Basset and DeepSEA, it achieved a lower mean squared error (MSE) than all other models (FigS4.a). Among multi-task models, DeepSEA outperformed all others across tissues and cell lines, while DanQ, SATORI, CNN+Attention, and Basset displayed comparable performance. In contrast, ExplaiNN, CNN+Transformer, and Scover (CNN) performed the worst (FigS4.b). Notably, aside from universally active chromatin accessible regions, sequence-based deep learning models did not exhibit substantial omissions of specific OCR categories (FigS3.c, FigS3.d). We also assessed the performance of two training strategies on both single-task and multi-task datasets. Unlike binary classification tasks, although each strategy performed well on its respective dataset, cross-dataset transfer tests revealed significant performance declines. Overall, multi-task learning models outperformed single-task models in regression tasks (Fig2.c, Fig2.d).

Models designed for long-sequence modeling, such as Basenji^23^ and Enformer, have demonstrated exceptional performance in predicting epigenomic marks across the whole genome. Comprehensive evaluations of these approaches have been conducted by S. Toneyan and colleagues. By visualizing genomic tracks of observed signals alongside model predictions, we found that Enformer produced smoother global predictions with closer alignment to experimental signals, attributable to its effective modeling of long-range interactions. In contrast, single-peak regression and classification methods exhibited higher false-positive rates in localized regions and were less accurate and consistent than Enformer (Fig2.e). However, single-peak methods offer advantages in terms of parameter efficiency, ease of training, and their utility in interpreting sequence-function relationships through explainable approaches.

### Optimal Data Processing Strategies and Model Robustness

Data preprocessing is a critical factor influencing both model performance and interpretability. Various data processing strategies have been adopted across models, including the selection of input sequence lengths and negative sample sampling methods. For instance, Basset uses sequences of 256 bp, DanQ and ExplaiNN utilize 600 bp, while DeepSEA and Scover employ 1000 bp. Dataset size also significantly impacts model performance. To identify optimal data processing strategies, we evaluated the effects of different input sequence lengths (200, 400, 600, 800, and 1000 bp) on model performance and compared the effectiveness of selecting negative samples from inactive regions versus generating synthetic negative samples. Additionally, we analyzed the impact of dataset size on model performance (Fig3.a). Based on the distribution of peak lengths, an input length of 200 bp appears appropriate, as most peaks fall within the range of 200–500 bp (FigS5.a). Training times for different models also varied with input sequence length (FigS5.b). Simpler models required shorter training times, whereas more complex models, such as SATORI, which integrates CNN, LSTM, and self-attention mechanisms, required the longest training times. For LSTM-based models, training time increased linearly with input sequence length, while non-LSTM models exhibited training times of approximately 100–150 seconds per epoch (batch size = 128) across sequence lengths ranging from 200 to 1000 bp.Most models demonstrated improved AUROC values with increasing input sequence length. Specifically, DeepSEA, Basset, SATORI, CNN, and ExplaiNN achieved their best performance at sequence lengths of 600 bp or 1000 bp (Fig3.b), suggesting these lengths are optimal for binary classification tasks. A similar trend was observed for regression tasks, where Pearson correlation coefficients (PCCs) improved with longer input sequences (Fig3.b).

**Fig 3.**
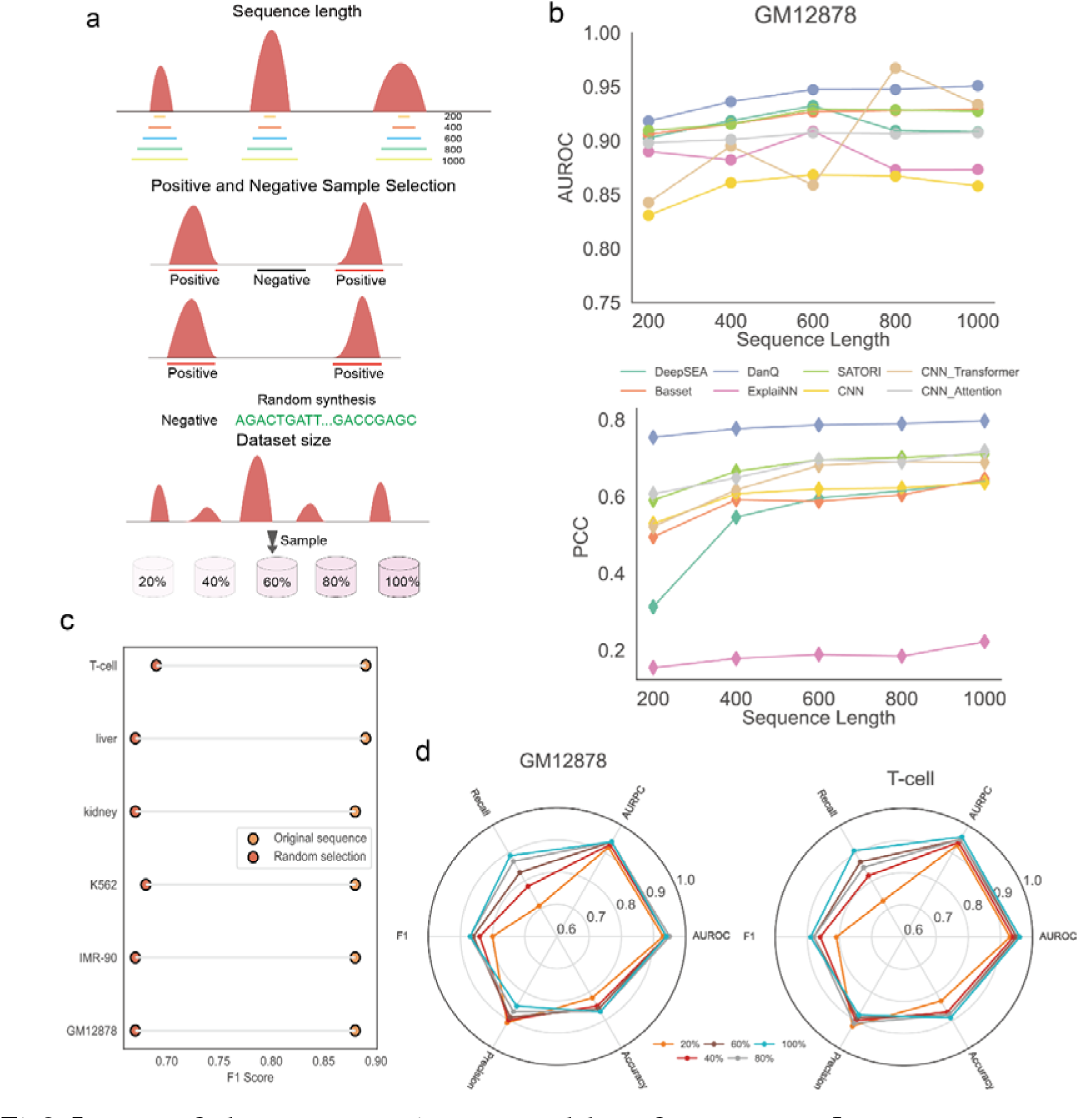
Impact of dataset processing on model performance. **a,** Input sequence lengths include 200, 400, 600, 800, and 1000 bp (top). Negative sample selection includes two approaches: from naturally inactive regions and synthetic sequences (middle). Different simulated data volumes include 20%, 40%, 60%, 80%, and 100% of the full dataset (bottom). **b**, The AUROC across different input sequence lengths on the GM12878 chromatin accessibility dataset (top). PCC across different input sequence lengths on the GM12878 chromatin accessibility dataset (bottom). **c**, Performance of negative sample selection from natural inactive regions and synthetic negative sample. d, Model performance with different dataset sizes for GM12878 and T-cells.

In binary classification tasks, we compared two methods for selecting negative samples: choosing non-active regions and generating random synthetic sequences. Across all six datasets, the approach of selecting non-active regions significantly outperformed the use of random synthetic sequences, with the former achieving F1 scores consistently above 0.85, while the latter remained below 0.7 (Fig3.c). This discrepancy is likely attributable to the pronounced genomic differences between non-active and active regions, which are absent in randomly generated sequences, thereby limiting the model’s ability to distinguish between classes. Dataset size also had a substantial impact on model performance. For instance, the GM12878 dataset contains 223,152 peaks with p-values below 0.05, while the liver and kidney datasets include approximately 180,000 peaks each. We evaluated model performance across datasets of varying sizes and observed that using 20% of the dataset yielded a recall of 0.60, whereas using 60% increased recall to over 0.8 (Fig3.d). Furthermore, model performance was strongly correlated with signal intensity, declining as signal values decreased (FigS5.c). Based on the comprehensive evaluation, we conclude that an input sequence length of 600 bp provides an optimal balance between training time and performance. Additionally, selecting non-active regions as negative samples represents the most effective strategy. While sequence-based deep learning models perform well even on datasets with limited cell numbers, we recommend acquiring as much data as possible to further enhance model performance.

### Performance of single-cell chromatin accessibility cell sequence representations

In single-cell ATAC-seq (scATAC-seq) data analysis, traditional methods typically rely on inter-cellular similarity for denoising and modeling, often neglecting the contribution of DNA sequences. Biological studies have demonstrated that chromatin accessibility is marked by transcription factor binding sites, which interact with specific DNA sequences. To address this limitation, Yuan et al. proposed scBasset^21^, a convolutional neural network (CNN)-based method for modeling scATAC-seq data. This approach predicts chromatin accessibility from DNA sequences and supports single-cell representation learning for downstream applications such as visualization and clustering. Although scBasset accurately predicts chromatin accessibility from sequences using deep CNNs, the performance of other sequence-based deep learning methods in scATAC-seq data analysis remains unexplored. To address this gap, we conducted a comprehensive evaluation of several models using the Buenrostro 2018 dataset^35^. Following preprocessing, we retained 2,034 cells and converted the data into a binary count matrix for each cell. A 1,344 bp DNA sequence centered on each peak was extracted, one-hot encoded, and used as input to eight models adapted for single-cell tasks. All models employed the same batch normalization, max-pooling, and Gaussian Error Linear Unit (GELU) activation functions as scBasset. To extract low-dimensional representations of peaks and learn cell-level features, we incorporated bottleneck layers into each model. The bottleneck embeddings were linearly transformed to predict binary accessibility for each cell (Fig4.a). All models were trained using binary cross-entropy loss and optimized with the Adam optimizer.

**Fig 4.**
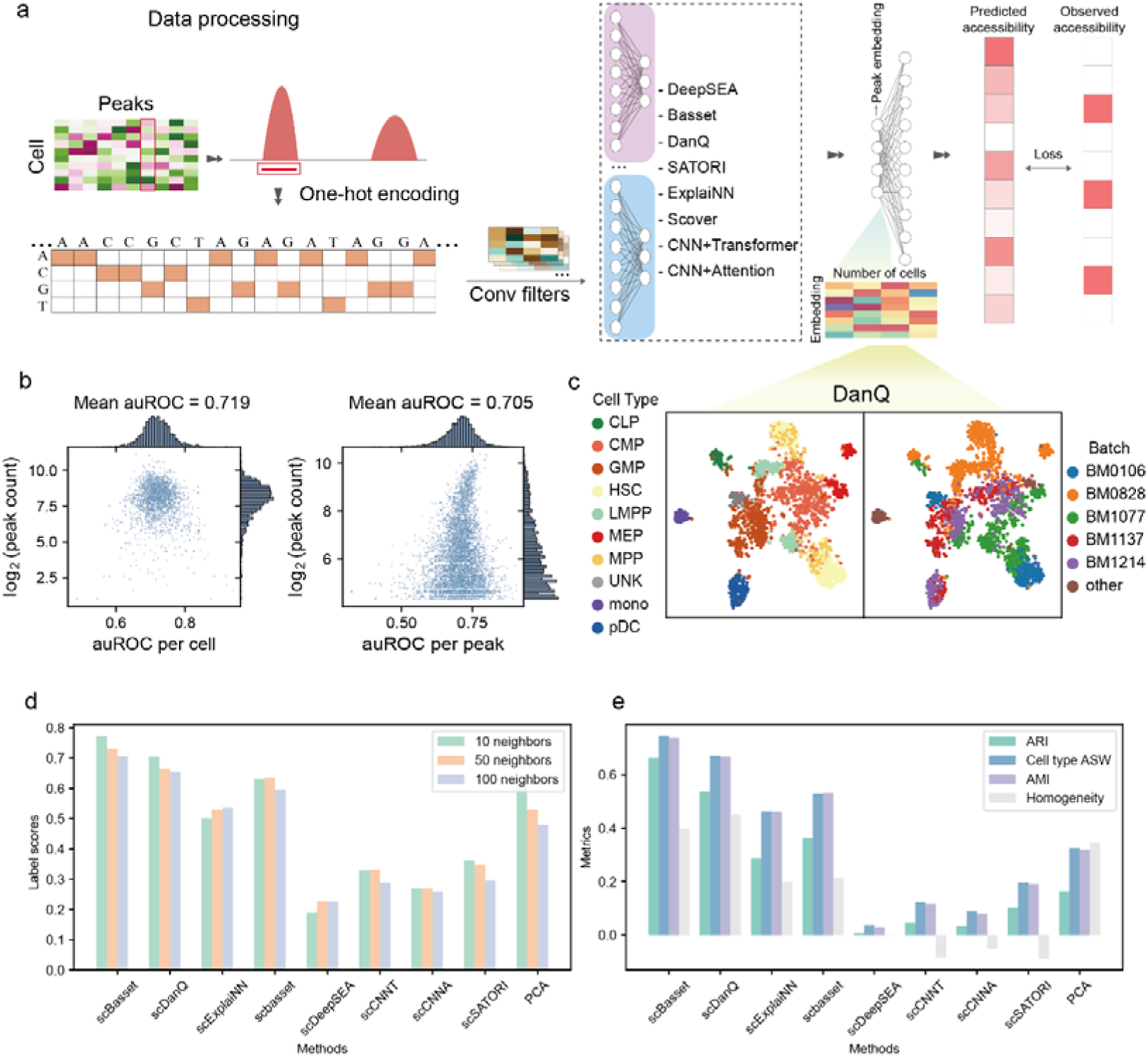
Evaluation of sequence-based models for chromatin accessibility prediction and representation learning in scATAC-seq data. **a,** Overview of the experimental pipeline for evaluating chromatin accessibility models on scATAC-seq data. **b**, Performance of DanQ in predicting chromatin accessibility at the single-cell and peak levels. **c**, t-SNE visualization of single-cell embeddings generated by scBasset and DanQ. **d**, Performance comparison of different cell-embedding methods evaluated by label score the proportion of cells’ nearest neighbors that share its cell type label—on buenrostro2018 dataset. **e**, Performance comparison of different cell-embedding methods evaluated by clustering metrics (ARI, cell type ASW and AMI) on the buenrostro2018 dataset.

We first evaluated the ability of various methods to predict chromatin accessibility for individual cells while preserving peak sequences, ensuring that the models could learn the relationship between DNA sequences and accessibility under sparse and noisy labeling conditions. For retained peaks, we calculated the area under the receiver operating characteristic curve (auROC) for each method at both the single-cell and peak levels. The results indicated that both DanQ and scBasset achieved robust performance: DanQ attained auROC scores of 0.719 per cell and 0.705 per peak, while scBasset achieved scores of 0.728 per cell and 0.707 per peak (Fig4.b, FigS6.a, FigS6.b). Apart from SATORI, other methods also demonstrated a certain degree of capability in predicting single-cell ATAC-seq accessibility.

Furthermore, the learned weight matrices were utilized to generate low-dimensional representations of single cells, which were compared against principal component analysis (PCA). Visualizing the cell embeddings using t-SNE (Fig4.c, FigS6.d, FigS6.e) revealed that scBasset and DanQ effectively clustered cells of the same type within the embedding space. Using Louvain clustering and comparing the results with true cell type labels, we computed the adjusted Rand index (ARI), adjusted mutual information (AMI), and homogeneity scores. scBasset and DanQ outperformed other methods across these metrics (Fig4.d, Fig4.e). Additionally, scBasset’s embeddings were used to construct a neighborhood graph and compute "label scores," demonstrating its superior ability to cluster cells of the same type compared to alternative approaches. Despite not being specifically optimized for single-cell ATAC-seq data, DanQ also exhibited strong performance, indicating the potential of its architecture for analyzing single-cell ATAC-seq datasets.

### Evaluating the interpretability and efficiency of models

A major downstream application of sequence-function models is interpretability analysis, which enables the identification of functional motifs within sequences and their complex interactions^36^. High predictive accuracy on held-out datasets suggests that these models implicitly learn key features of the regulatory code^37^.

Interpretability methods for sequence-function models can be broadly categorized into attribution map-based methods (AMBM) and sequence alignment-based methods (SABM) (Fig5.a). AMBM approaches, such as Input × Gradient, DeepLIFT, and Saliency Map, compute importance scores (IS) for each position in the input sequence via gradient backpropagation and highlight critical regions through visualization^38, 39^. Tools like TF-MoDISco further refine this process by clustering importance scores to estimate position weight matrices (PWMs)^40^. In contrast, SABM approaches identify subsequences that activate convolutional neurons and stack PWMs, analogous to traditional estimation methods^41^. While convolutional neural networks (CNNs) have been widely employed to identify cell type-specific sequence syntax, differences in performance between models and their ability to learn functional motifs remain unclear. Here, we systematically compared several commonly used interpretability methods on binary and quantitative models, focusing on functional motif discovery and the identification of important sequence regions.

**Fig 5.**
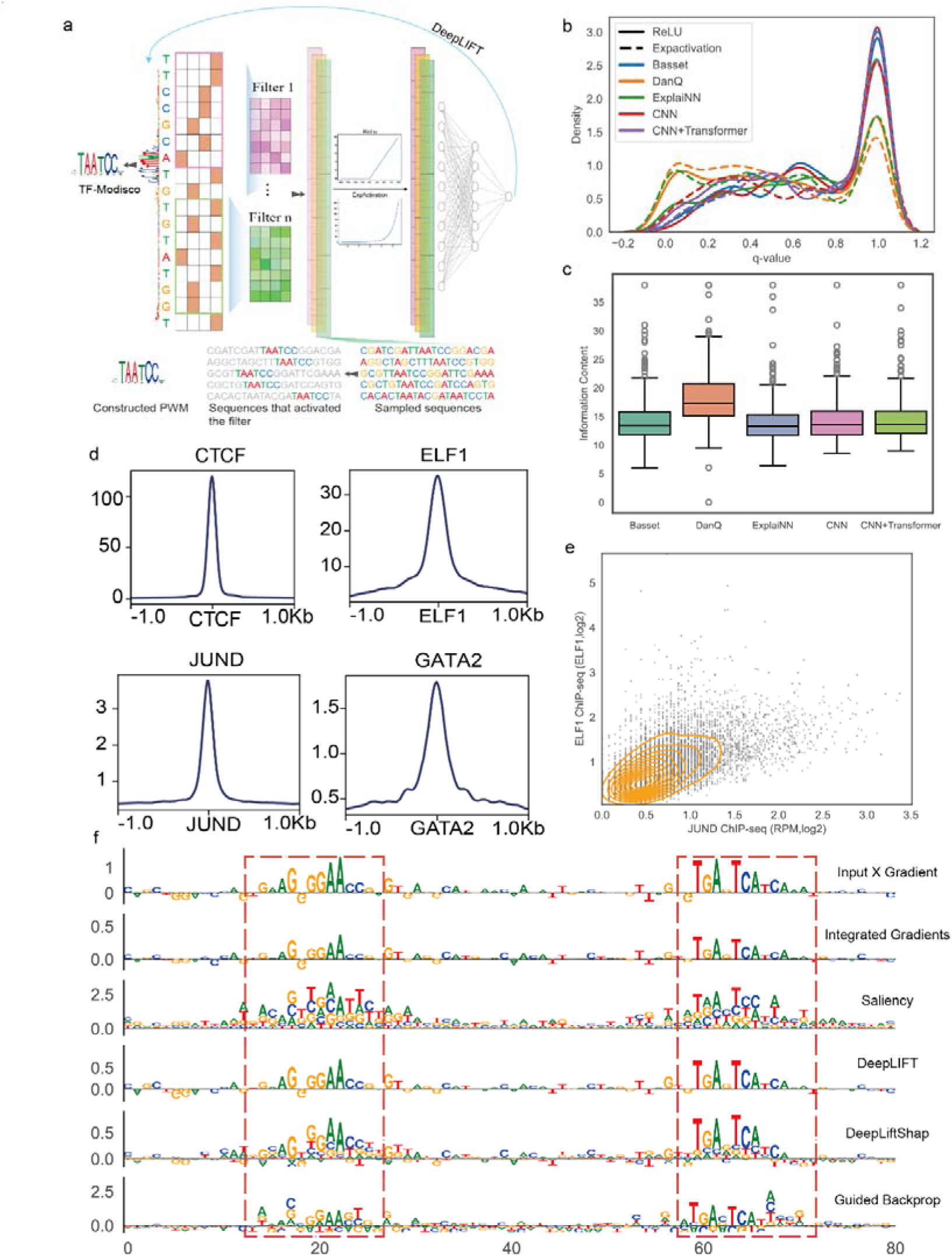
Model interpretability in learning sequence grammar. **a,** Interpretable methods for sequence-function modelling: sequence alignment based method (SABM) and attribution map based method (AMBM). **b,** The q-value generated by the motifs found by the different models after TOMTOM comparison with known motifs in Jasspar database. **c,** IC values (using the ReLU activation function) across different methods. **d,** Significant regions identified by model and ChIP-seq for known transcription factors overlap. **e,** Transcription factor synergies found by the model and synergies found by ChIP-seq experiments. **f,** Different attribution methods yield distinct attribution scores at the AP-1 motif.

We employed q-value (minimum false discovery rate) and information content (IC) to evaluate each model’s ability to identify functional motifs. A smaller q-value indicates lower false-positive similarity between the discovered motifs and known motifs in databases^42, 43^. Previous studies have shown that the choice of activation function influences a model’s capacity to learn sequence motifs^44^. Therefore, we assessed the effect of ReLU and exponential activation functions on model interpretability. DanQ consistently achieved the lowest q-values and the highest IC regardless of whether ReLU or the exponential activation function was used, reflecting its overall superior performance (Fig5.b, Fig5.c). We observed that the exponential activation function generally outperformed ReLU. ExplaiNN, a model specifically designed for interpretability, also achieved competitive q-values when using the exponential activation function, comparable to DanQ with ReLU (Fig5.b, FigS7.a). However, other methods performed less favorably under both activation functions compared to DanQ and ExplaiNN.

To explore the relationship between input sequence length and model interpretability, we evaluated the q-values of DanQ across varying input lengths. The results were optimal at an input length of 200 bp (FigS8.a). Although model performance showed slight improvement at 1000 bp, its interpretability did not improve; in fact, it was inferior to the results obtained at 200 bp or 600 bp. Quantifying the importance of individual filters represents another critical aspect of the SABM interpretability framework, as this metric can elucidate the contribution of distinct functional motifs to model predictions. We also assessed the efficiency of different models. The recurrent architecture of DanQ incurred the longest runtime, followed by Basset, which exhibited slower performance due to its extensive use of convolutional filters and linear neurons. In contrast, ExplaiNN, designed specifically for interpretability, demonstrated near-zero runtime for filter importance quantification (FigS8.b).

Attribution map-based methods (AMBM) quantify the contribution of individual nucleotides to model predictions, facilitating the identification of minimal functional elements. To enhance model interpretability, we propose a hybrid architecture that combines DanQ or Basset with ExplaiNN, leveraging the predictive accuracy of DanQ or Basset and the interpretability of ExplaiNN. We employed DeepLIFT to evaluate the ability of different methods to identify important regions and further assessed various attribution approaches on the Basset+ExplaiNN hybrid model (Fig5.f, FigS8.f). The results demonstrated that both Basset and Basset+ExplaiNN could more clearly identify key genomic regions. Among the attribution methods tested, Input × Gradient and DeepLIFT exhibited superior clarity in highlighting important regions. Additionally, motifs clustered using TF-MoDISco showed strong consistency with known motifs (FigS9.a, FigS9.c), whereas de novo motifs identified through MEME enrichment showed lower similarity to known motifs (FigS9.b).

By perturbing these important sites, we observed that the number of critical regions within a sequence significantly influences the robustness of model predictions. When sequences contained only 1–3 critical regions, perturbing the most important region led to a performance drop exceeding 70%. In contrast, sequences with a larger number of critical regions exhibited a smaller decrease of approximately 40% (FigS8.c, FigS8.d). This effect may arise from the redundancy of multiple transcription factor binding sites, where distinct binding sites exhibit complementary roles in gene regulation. We further validated the identified key regulatory regions using ChIP-seq data and found significant overlap between these regions and the experimental data (Fig5.d). To explore whether the co-occurrence of motifs for two different transcription factors within a sequence reflects cooperative interactions, we extracted sequences containing AP-1 and ELF1 motifs and visualized the read counts for the two transcription factors in these regions. The ChIP-seq signals of AP-1 and ELF1 showed a linear relationship, indicating that sequence-based methods can effectively capture cooperative effects and infer transcription factor binding patterns without the need for experimental data (Fig5.e).

In the study of non-coding variants, we analyzed 10,986 autoimmune-associated GWAS SNPs using the PICS statistical method. We observed that SNPs located in regions of high importance exhibited higher PICS scores, suggesting that these variants are more likely to influence chromatin accessibility and drive disease (FigS8.e). Attribution methods not only identify key regulatory elements and quantify their importance but also prioritize the functional effects of non-coding variants, providing a powerful tool for interpreting disease-associated loci.

### Cross-species conservation analysis of chromatin accessibility determinants via deep learning

The functional study of cis-regulatory elements (CREs) in fetal development and disease mechanisms often relies on model organisms^2, 45^. While mutational analyses of conserved orthologous elements in model animals have validated some hypotheses regarding human CRE functions, the rapid evolutionary divergence of CREs, such as enhancers, poses significant challenges to identifying homologous elements across species^45–47^. Sequence alignment-based approaches have been employed to construct cross-species enhancer maps but are limited in fully elucidating regulatory logic^8, 48^. Deep learning models, with their ability to learn regulatory syntax, offer a powerful tool for systematically studying the conservation and diversity of enhancers across species.

Here, we trained models using chromatin accessibility from early embryonic development in vertebrates, including humans, mice, cattle, chickens, medaka, and zebrafish^49, 50^. The models achieved high accuracy in predicting chromatin accessibility during early embryonic stages in humans and mice (AUROC: 0.919 for the human 8-cell stage; 0.888 for the mouse 2-cell stage; Fig6.a). Using sequence alignment-based methods (SABM), we successfully extracted the various sequence motifs learned by the model and identified that motifs associated with the OBOX family of transcription factors play a crucial role in predicting chromatin accessibility during the early stages of embryonic development. This finding aligns with current research, suggesting that OBOX transcription factors play a key role in regulating genome activation in the mouse zygote and in the early stages of embryogenesis (FigS10.b, FigS10.c)^51^. Species-specific models trained on datasets from other species also demonstrated high predictive accuracy (FigS11.a). These findings indicate that sequence-based deep learning models effectively capture key relationships between sequences and chromatin accessibility across diverse species.

Using LiftOver mapping^52, 53^, we identified conserved open chromatin regions across different species, accounting for 2.1% to 18.1% of the total open chromatin regions (Fig6.b). Further analysis revealed that the sequence determinants of chromatin accessibility are conserved across developmental stages within the same species but exhibit species-specific characteristics between humans and mice (Fig6.c). Despite the limited conservation of open chromatin regions between humans and zebrafish, their sequence determinants showed a high degree of concordance (FigS10.a). Additionally, the predictive scores for conserved regions were significantly higher than those for non-accessible regions, suggesting a degree of conservation in sequence determinants across species (Fig6.d). Among different types of CREs, sequence determinants of promoters and short interspersed nuclear elements (SINEs) were significantly more conserved than those of other types, supporting the cross-species conservation of promoter regions. In contrast, enhancers displayed a higher evolutionary rate (Fig6.e). Analysis of H3K27ac data revealed that regions with high predictive scores in cross-species models exhibited greater enhancer activity compared to regions with low predictive scores or LiftOver-aligned regions, a trend consistently observed in both human and mouse models (Fig6.e, Fig6.g). These findings demonstrate that sequence-based deep learning methods effectively uncover conserved sequence determinants of chromatin accessibility, providing critical insights into the conservation of enhancers across species.

**Fig 6.**
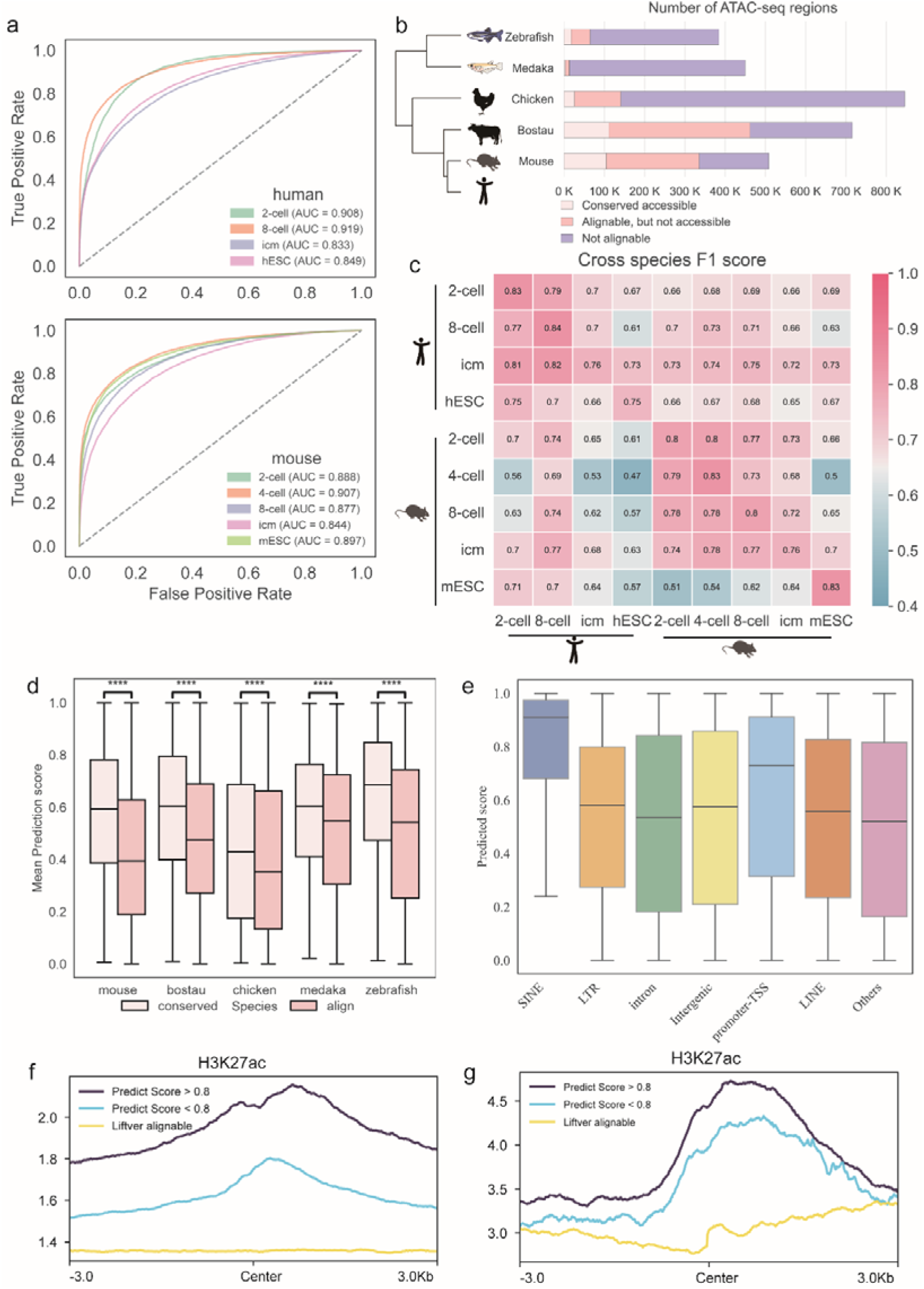
Cross-species enhancer logic analysis using sequence-function modelling. **a** Model performance on chromatin accessibility data during human and mouse early embryonic development. **b** The total number of ATAC-seq regions observed in all samples of a species is coloured according to whether they are non-alignable, alignable or conservatively accessible in humans. **c** Human and mouse cross-species chromatin accessibility prediction F1 score. **d** Predicted scores for the align regions and the conservative accessible regions. **e** Predictive scores of human models across species on different regulatory elements in mouse. **f** Comparison of enhancer activity in high and low prediction score regions identified by the mouse model on the human dataset, and regions identified through LiftOver. **g** Comparison of enhancer activity in high and low prediction score regions identified by the human model on the mouse dataset, and regions identified through LiftOver.

### Guidelines for optimal model selection

Through a series of benchmark tests, we have identified the optimal approaches for various scenarios. Taking into account the performance of each method and the characteristics of their associated architectures, we provide specific recommendations and guidelines tailored to common downstream applications. For tasks aimed at predicting genome-wide chromatin activity or quantifying chromatin accessibility sequencing quality using sequence-based deep learning methods, we recommend using an input sequence length of 600 bp to balance accuracy and training time. We suggest selecting negative samples from naturally inactive regions rather than artificially synthesized sequences, and we recommend using the ReLU activation function. Suitable model choices include DanQ or SATORI for binary, DeepSEA or Basset for regression. When learning motifs from sequences using chromatin accessibility data, we recommend using an input sequence length of 200 or 600 bp with an exponential activation function. Models such as DanQ, ExplaiNN, and Basset are recommended, and SABM interpretability methods can be used to extract sequence motifs from the models. For tasks focused on predicting the impact of sequence motifs on chromatin accessibility, we recommend using an input sequence length of 200 or 600 bp with an exponential activation function and selecting a model like ExplaiNN or Hybrid model, which is linearly additive. The use of SABM interpretability methods will facilitate the quantification of motif effects. When the task is to predict the effects of non-coding variants, we recommend using the more efficient AMBM method to quantify the impact of each nucleotide, allowing researchers to prioritize non-coding SNPs. For cross-species enhancer logic analysis, we recommend using a 600 bp sequence as input. A hybrid model can ensure both model performance and the quick extraction and quantification of sequence motifs. For the analysis of single-cell ATAC-seq data, we recommend employing the scBasset or DanQ models, as they demonstrate superior performance in learning cellular representations compared to other models. In summary, we hope the provided software package and recommendations will accelerate the development of sequence-based deep learning methods and related interpretability techniques. Most importantly, we aim to foster the development of more applications that leverage AI in functional genomics research.

**Fig 7.**
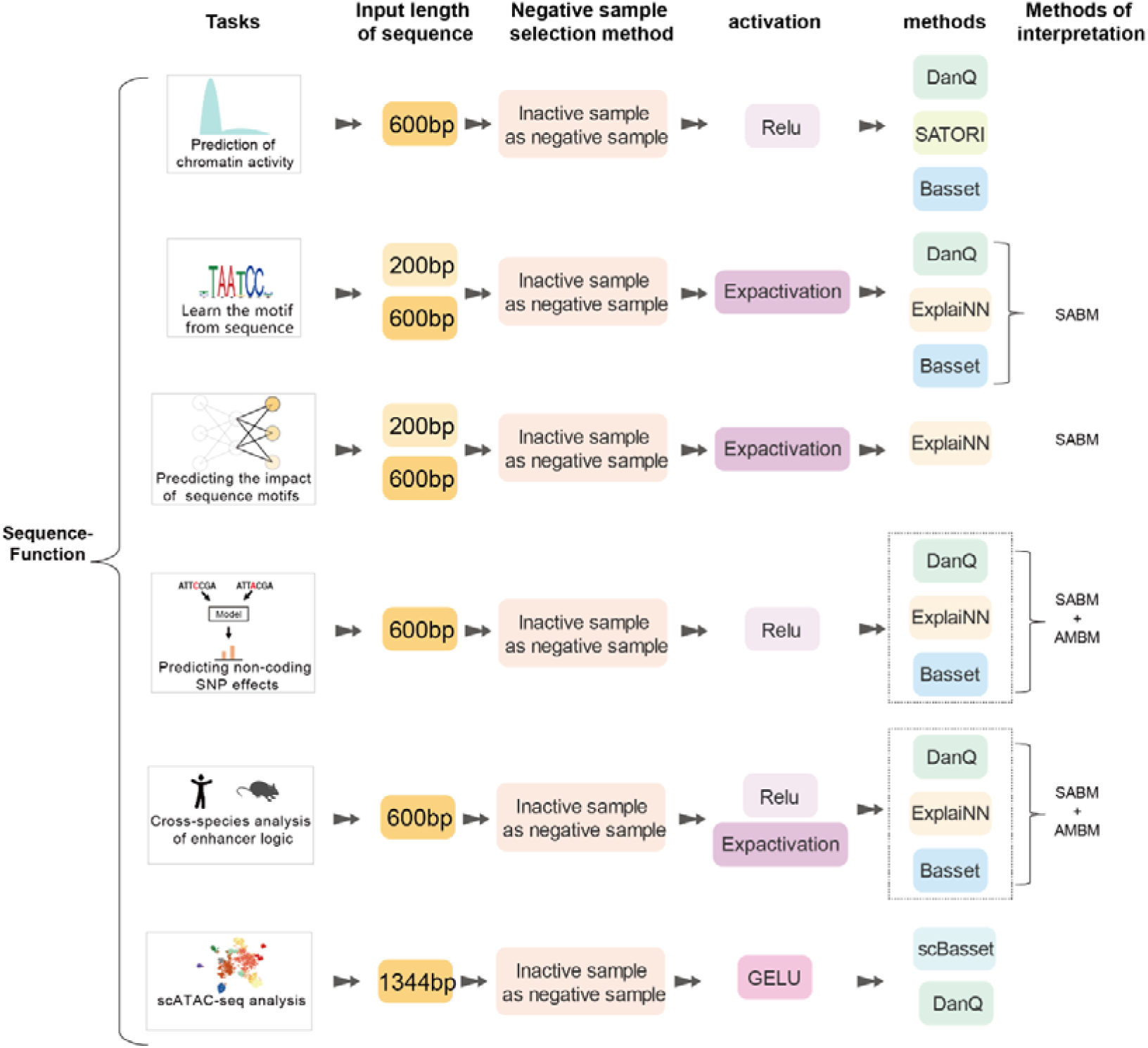
Scenario-specific guidelines for users. Six common scenarios are included, and the best option recommends for each scenario.

## Discussion

In this study, we proposed a comprehensive evaluation framework to systematically assess sequence-based chromatin accessibility prediction methods across 369 cell lines and tissues. Our findings demonstrate that LSTM-based and multilayer convolutional model achieved better prediction accuracy in binary classification and quantitative regression tasks. Models based on self-attention mechanisms did not show outstanding performance, despite their success in natural language processing and image processing. Simpler model architectures, such as Scover, which is based on a single-layer convolutional neural network and a single-layer linear neural network, exhibited the weakest performance, likely due to their inability to effectively extract the sequence rules necessary for classification or regression tasks.

Compared to existing benchmark studies, our research focuses on methods that predict single peak signals from DNA sequences, rather than long-sequence chromatin signal prediction methods like Basenji and Enformer. Long-sequence input methods can accurately predict epigenomic signals by integrating interactions between regulatory elements within the sequence for final predictions. However, interpreting these models and extracting biologically meaningful insights is challenging due to the large number of parameters involved. Although recent studies have employed AMBM to interpret these models and investigate interactions between cis-regulatory elements, these models still lack interpretability in specific scenarios compared to binary classification or quantitative regression models trained on single peaks. This study encompasses the majority of currently available methods for predicting chromatin accessibility within individual peaks. Together with the work of *Toneyan, S. et al*., it provides a comprehensive overview of sequence-function models, offering valuable insights into this domain^54^.

Based on our evaluation, we believe that there are several common but critical issues in current sequence-based functional genomics models that remain unresolved. First, although we demonstrated that using naturally inactive regions as negative samples is a good choice, these regions may contain false negatives, a problem that could be alleviated with advancements in sequencing technology. Second, model interpretability methods still need improvement. While SABM can extract sequence rules from models, most of the motifs learned by the filters cannot be matched to motifs in existing databases. This discrepancy may be due to the discovery of novel motifs or the limitations of current interpretability methods. Recent research addressing the problem of multifunctional neurons in sequence-function models has optimized neuron representations by clustering the subsequences that activate them into different groups^29^. Finally, while training with a reference genome may pose a challenge in quantifying non-coding SNPs, the inclusion of paired individual genomic and epigenomic data can alleviate this problem^55^.

Potential directions for future development in this area may also include the following: 1) The accumulation of single-cell epigenomics data provides opportunities to study the regulatory programs of cellular heterogeneity using deep learning, with the hope that more methods and tools suitable for single-cell cis-regulatory analysis will be developed. 2) The generation of personalized genome-paired omics data enables the training of more individual-specific sequence-function models, aiding in the investigation of the relationships between specific SNPs and particular phenotypes and diseases. 3) The development of transparent models and interpretability methods. Many algorithms for interpreting complex models have already been developed in natural language processing and image processing. Future research should focus on successfully applying these methods to functional genomics research.

## Methods

### Data collection and preprocessing

The ATAC-seq data were obtained from the Encyclopedia of DNA Elements (ENCODE) database. The bigWig tracks were log-fold normalized for sequencing depth using control tracks, following the data processing pipeline established by ENCODE. No further processing was performed. The ATAC-seq datasets used in the analysis correspond to the following ENCODE experiment numbers: ENCFF470YYO (GM12878), ENCFF273DGN (T-cell), ENCFF243NTP (IMR-90), ENCFF842UZU (K562), ENCFF845OZO (Kidney), and ENCFF488BRH (Liver). The reference genome used for all datasets was hg38. The chromatin accessibility data during early vertebrate embryonic development were derived from our previous work, including samples from human (2-cell, 8-cell, ICM, hESC), mouse (2-cell, 4-cell, 8-cell, ICM, mESC), bostau (2-cell, 4-cell, 8-cell, ICM, morula, ESC), chicken (HH11, HH16, HH19, HH24, HH28, HH32, HH38, HH6), medaka (st15, st21, st28, st36, st24, st32, st40), and zebrafish (1k-cell, 256-cell, 64-cell, dome, oblong). The reference genomes used were hg19 for human, mm10 for mouse, Bostau9 for cattle, Galgal5 for chicken, OryLat2 for medaka, and DanRer10 for zebrafish. The human data were obtained from GSE101571. The mouse data were obtained from GSE66581, GSE66582, and DRP004498. The bovine data were obtained from GSE143658 and GSE52415. The chicken and medaka data were obtained from DRP004498. The zebrafish data were obtained from GSE101779, GSE106428, GSM1289382, and GSE106431.

### Positive and negative training samples

For binary classification tasks, we defined positive samples by taking the midpoint of each peak as the center and extracting DNA sequences of length L/2 on both flanking sides, where L is the input sequence length. Non-overlapping, inactive regions were selected as negative samples. For synthetic negative samples, we generated DNA sequences randomly using a common approach, with each of the four bases having an equal probability of 0.25. In regression tasks, we used pyBigWig to extract normalized coverage values from BigWig files.

### Data splits

We split the dataset into training and test sets, with chromosomes 8 and 9 designated as the test set and the remaining chromosomes used for training. Additionally, we removed all uninformative regions from the segmented data. The dataset was split consistently across different experiments to enable direct comparison.

## Models

### DeepSEA

The DeepSEA model is composed of a convolutional block followed by two max-pooling layers and a fully connected layer. The first convolutional layer utilizes a kernel size of 8 and 320 filters to extract initial features from the input DNA sequences. This is followed by a nonlinear transformation through an activation function. The model then applies a max-pooling operation with a pooling size of 2, reducing the resolution of the feature map and computational load. The second convolutional layer employs 480 filters with the same kernel size, processed again by a ReLU activation function and followed by max-pooling. The third convolutional layer uses 960 filters, introduces batch normalization, and then applies ReLU activation and dropout to mitigate overfitting. After the convolutional layers, the model proceeds to a fully connected layer, where the features are reduced to 256 units, activated by ReLU, and regularized with dropout, ultimately leading to a dense layer that outputs the classification results.

### Basset

The Basset model is also based on a convolutional neural network, consisting of multiple convolutional layers and max-pooling layers, followed by a fully connected layer. The first convolutional layer uses a kernel size of 11 and 300 filters, with batch normalization and an activation function to process the input sequences. This is followed by max-pooling with a pooling size of 3 and a stride of 2. The second convolutional layer uses a kernel size of 11 and 200 filters, followed by batch normalization, ReLU activation, and max-pooling with a pooling size of 4 and a stride of 2. The third convolutional layer uses a kernel size of 7 and 200 filters, once again followed by batch normalization, activation, and max-pooling. The fully connected layer is composed of three stages, first reducing the features to 1000 units, processed by ReLU activation and regularized with dropout multiple times, leading to the final dense layer that outputs the prediction.

### DanQ

The DanQ model combines a convolutional neural network (CNN) with a long short-term memory network (LSTM). The model begins with a convolutional layer that includes 320 filters and a kernel size of 26, processing the input sequences. This is followed by max-pooling with a pooling size of 13 and a stride of 13, along with dropout to reduce overfitting. The feature map generated by the convolutional layers is then fed into a bidirectional LSTM layer, which has 320 hidden units and consists of 2 LSTM layers, designed to learn long-range dependencies within the sequences. Finally, the model uses a fully connected layer to produce the predictions, with the preceding features processed by ReLU activation and further regularized with dropout to mitigate overfitting.

### ExplaiNN

The ExplaiNN model focuses on interpretability, starting with a convolutional layer that has 300 filters and a kernel size of 19 to process the input sequences. This layer is followed by batch normalization and an activation function. Max-pooling with a pooling size of 10 is then applied. After feature extraction, the model employs linear layers and sequential convolutional operations, utilizing grouped convolutions to enhance interpretability. The final output is generated through a fully connected layer that performs the classification.

### SATORI

The SATORI model integrates convolutional neural networks (CNN), long short-term memory networks (LSTM), and multi-head self-attention mechanisms. The first convolutional layer consists of 512 filters and a kernel size of 21, performing initial feature extraction, followed by max-pooling with a pooling size of 10 and dropout to reduce overfitting. The features are then passed to a bidirectional LSTM layer, with each layer containing 256 hidden units, to capture long-range dependencies in the sequence. Afterward, a multi-head self-attention mechanism is used to capture the interactions between features within the sequence. The final predictions are generated through a fully connected layer.

### CNN + Transformer

The CNN_Transformer model combines convolutional neural networks with the Transformer architecture. The input sequences are initially processed by a convolutional layer with 300 filters and a kernel size of 19. This is followed by max-pooling with a pooling size of 10 and dropout to reduce overfitting. The feature map produced by the convolutional layers is then input into the Transformer encoder layer, which captures long-range dependencies within the sequences. The final classification output is generated through a fully connected layer.

### CNN + Attention

The CNN_Attention model combines convolutional neural networks with multi-head self-attention mechanisms. The input sequences first pass through a convolutional layer with 300 filters and a kernel size of 19, followed by max-pooling with a pooling size of 10 and activation functions for further processing. The resulting convolutional feature map is then fed into a multi-head self-attention layer to capture interactions between features within the sequence. The final predictions are produced through a fully connected layer.

### CNN (Scover)

The CNN model is a basic convolutional neural network architecture. The input sequences are initially processed by a convolutional layer with 300 filters and a kernel size of 19, followed by batch normalization and an activation function. Max-pooling is then applied with a pooling size of 6 and a stride of 6. The convolutional feature map is passed to the fully connected layer, where it is first processed by ReLU activation and then regularized with dropout to reduce overfitting, finally producing the classification output through a dense layer.

### Hybrid Model

To achieve both accuracy and interpretability, we propose using DanQ or Basset as the backbone network, while incorporating grouped convolutions from the ExplaiNN model as side branches. The final output of the network integrates the results from both the backbone and the side branches. This hybrid model balances high accuracy with strong interpretability, making it highly adaptable for downstream applications.

### Training

For the binary classification models, we used the binary cross entropy loss function.

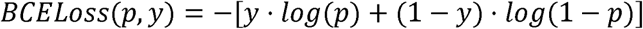

Where *p* represents the predicted probability (output of the model) that the label is 1. *Y* represents the actual label, where *y* ∈{0,1}. For regression models, we employed mean squared error and Pearson correlation coefficient as loss functions.

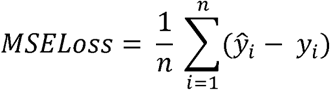

Where *ŷ_i_* represents the predicted value for the *i*-th sample, *y_i_* represents the actual target value for the *i*-th sample and *n* is the total number of samples in the batch. The Pearson loss function implemented in the provided code is designed to minimize the dissimilarity between the predicted and target vectors by penalizing the differences in their Pearson correlation. This loss is computed in several steps:

### Mean Centering

The input vectors *x* and *y* are first mean-centered by subtracting their respective means *u_x_* and *u_y_*. This step results in centered vectors 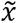 and 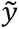, which are defined as:

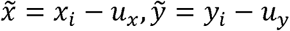

Where 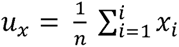 and 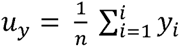.

### Cosine Similarity Calculation

The cosine similarity between the centered vectors 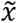 and 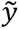 is then computed as:

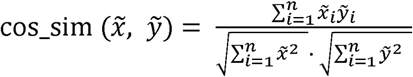

### Pearson Loss Computation

Finally, the Pearson loss is computed by taking the mean of 1− cos_sim 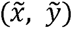 across all samples in the batch:

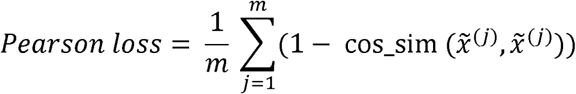

where mmm is the number of samples in the batch. This formulation encourages the model to minimize the angular difference between the predicted and target vectors, effectively promoting a higher Pearson correlation and thus a more accurate prediction.

Each model was trained for up to 100 epochs using the Adam optimizer default parameters. An early stopping strategy was implemented, with a patience set to 5 epochs. Due to the imbalance between positive and negative samples, we employed a sampling strategy to ensure that each batch had a balanced distribution of positive and negative samples. The initial learning rate was set at 0.001, and a dynamic learning rate schedule was used, where the learning rate was reduced by a factor of 0.2 every three steps.

### Methods comparison

We conducted a detailed comparison of the performance of eight different model architectures in predicting chromatin accessibility. Here, we provide an in-depth description of the model architectures and hyperparameter settings for these methods. To ensure a fair comparison, we standardized the input sequence length to 600 bp across all models. Additionally, we maintained consistent hyperparameter settings throughout the training process. To minimize errors associated with individual training runs, each model was trained ten times using different random seeds.

### scATAC-seq preprocessing

We utilized the processed peak set generated by Chen et al. and employed in the scBasset study, derived from Buenrostro2018. The dataset, accessible at https://github.com/pinellolab/scATAC-benchmarking/blob/master/Real_Data/Buenrostro_2018/input/combined.sorted.merged.bed, consists of peaks called from aggregated profiles of each cell type and subsequently merged into a unified atlas. The aligned BAM files, also provided by Chen et al., were downloaded from https://github.com/pinellolab/scATAC-benchmarking/tree/master/Real_Data/Buenrostro_2018/input/sc-bams_nodup. Peaks accessible in fewer than 1% of cells were excluded from the analysis. The final dataset comprised 103,151 peaks and 2,034 cells.

### scBasset model architecture

scBasset predicts binary accessibility for DNA peaks using a 1,344-bp sequence encoded as a 1,344×4 matrix. The architecture begins with a 1D convolutional layer containing 288 filters (kernel size 17) with batch normalization, GELU activation, and max-pooling, producing a 488×288 output. This is followed by a convolution tower with six blocks, where the number of filters increases from 288 to 512 (kernel size 5), combined with batch normalization, GELU, and max-pooling, resulting in a 7×512 matrix. A 1D convolutional layer with 256 filters (kernel size 1) and GELU generates a 7×256 matrix, which is flattened into a 1×1,792 vector. A dense bottleneck layer with 32 units, incorporating batch normalization, dropout (rate 0.2), and GELU, compresses the representation into a 1×32 vector. Finally, a dense layer predicts continuous accessibility scores.

### Analysis of single-cell ATAC-seq data using alternative methods

For the other methods, a bottleneck layer was introduced prior to the classification layer to adapt to the specific requirements of single-cell ATAC-seq data analysis.

### Cell-embedding evaluation

The weights of the bottleneck layer and the final classification layer from the model were utilized to derive low-dimensional embeddings for the cells.

### Clustering-based metrics

The quality of the learned cell embeddings was assessed by comparing the resulting clusters to the ground-truth labels: FACS-sorted cell type labels for the Buenrostro2018 dataset and RNA-based cell cluster labels for the multiome dataset.

### Cell Type Average Silhouette Width (ASW)

The silhouette width assesses the coherence of cell embeddings by measuring the relative proximity of a cell to others with the same label compared to those with different labels. To evaluate the quality of cell embeddings, we calculated the ASW, as proposed in previous single-cell studies, which represents the average silhouette score across all cells, re-normalized to a scale of 0 to 1.

### Label Score

The quality of the learned cell embeddings was assessed using the label score across all three datasets. The label score measures the percentage of a cell’s neighbors within a defined neighborhood that share the same label in the nearest-neighbor graph. For each embedding method, label scores were calculated for neighborhood sizes of 10, 50, and 100. As the ground-truth cell types for the multiome datasets are unavailable, we used cluster identifiers derived from scRNA-seq Leiden clustering as proxy cell-type labels.

## Model interpretability

### Sequence Alignment Based Method (SABM)

SABM is a feature map-based method used to interpret model parameters. This approach begins by extracting the filters from the first convolutional layer of the network and converting them into Position Weight Matrices (PWMs) by statistically counting the occurrences of deoxyribonucleotides in the feature maps. We set the activation threshold at 0.85, meaning that only nucleotides that activate the filter by at least 85% of its maximum value are included in the PWM. Subsequently, the TOMTOM algorithm is employed to query the JASPAR database, annotating the filters represented by the PWMs with known transcription factor binding sites (TFBS). As a result, corresponding transcription factor motifs are assigned to each filter.

### Filter influence

For each model, except ExplaiNN, we assessed the impact of each filter by replacing all activation values with zero tensors and comparing the fold change between the modified and original predictions. This fold change was used to quantify the filter’s influence. For ExplaiNN, which utilizes the concept of a generalized linear additive model, group convolution is employed to prevent interactions between feature maps of different filters, and a linear model is then used to fit the filter outputs. Consequently, the weights of this final linear model can be used to quantify the influence of each filter.

### Attribution Map Based Methods (AMBM)

AMBM interprets model parameters using attribution analysis. Common feature attribution methods, such as Input x Gradient, DeepLIFT, and Saliency Map, utilize gradient backpropagation algorithms to obtain importance scores (IS) for each position in the input sequence and visualize portions of the IS to highlight critical sequence regions. We employed the Captum package to execute the Input x Gradient algorithm and visualized the attribution maps using TF-MoDISco. Finally, we used TF-MoDISco to cluster the attribution results into Position Weight Matrices (PWMs).

### Identification of attribution-important regions

We scanned the entire sequence’s attribution values using a sliding window of width *d* and employed the *find_peaks* function from the *scipy* package in Python to identify regions with high attribution scores. Regions were selected using a threshold set at 0.3 of the peak maximum, with a minimum distance of *d* between regions, corresponding to the size of the sliding window.

### PICS

We followed the approach provided by Basset and downloaded the PICS SNP annotations from the supplementary materials of the study by Farh et al. (2015). We then filtered out all SNPs associated with protein-coding genes, focusing exclusively on the non-coding SNP set.

### Defining sets of alignable and conserved accessible ATAC-seq regions

If the ATAC-seq regions from non-human species can be converted to hg19 coordinates using the liftOver tool with appropriate liftOver chains (UCSC), they are defined as alignable regions. We intersected these alignable regions with accessible peaks in humans using the –element-of function in bedops (v2.4.41) and applied the -n 1 option to define the set of conserved accessible regions across species.

## Data availability

In this study, we collected data from public databases. ATAC-seq, H3K27ac, and ChIP-seq data were obtained from the ENCODE project for GM12878, T-cell Liver, Kidney, IMR-90 and K562 cell lines, and from Gene Expression Omnibus (GEO) with accession number GSE101571 for Human. The mouse data were obtained from GSE66581, GSE66582 and DRP004498. The bovine data were obtained from GSE143658 and GSE52415. The chicken and medaka data were obtained from DRP004498, in which the whole embryo was used to perform ATAC-seq.

## Software availability

Source code implementing all steps—data preprocessing, training, and downstream analysis—is available in the package cisFinder from https://github.com/bioczsun/cisFinder.

## Declarations

### Author contributions

H.C., M.L., and X.B. conceived and supervised the project. C.S., Y.S. and K.X. designed the framework and performed data analysis with help from Z.H., H.L., Y.L., Z.Y., Y.W., X.L., X.X., P.H. H.C., M.L., X.B. H.C., M.L., X.B. and C.S drafted the manuscript. All authors read and approved the final manuscript.

