## supplementary information 1 for "A comprehensive benchmark and guide for sequence-function interpretable deep learning models in genomics"

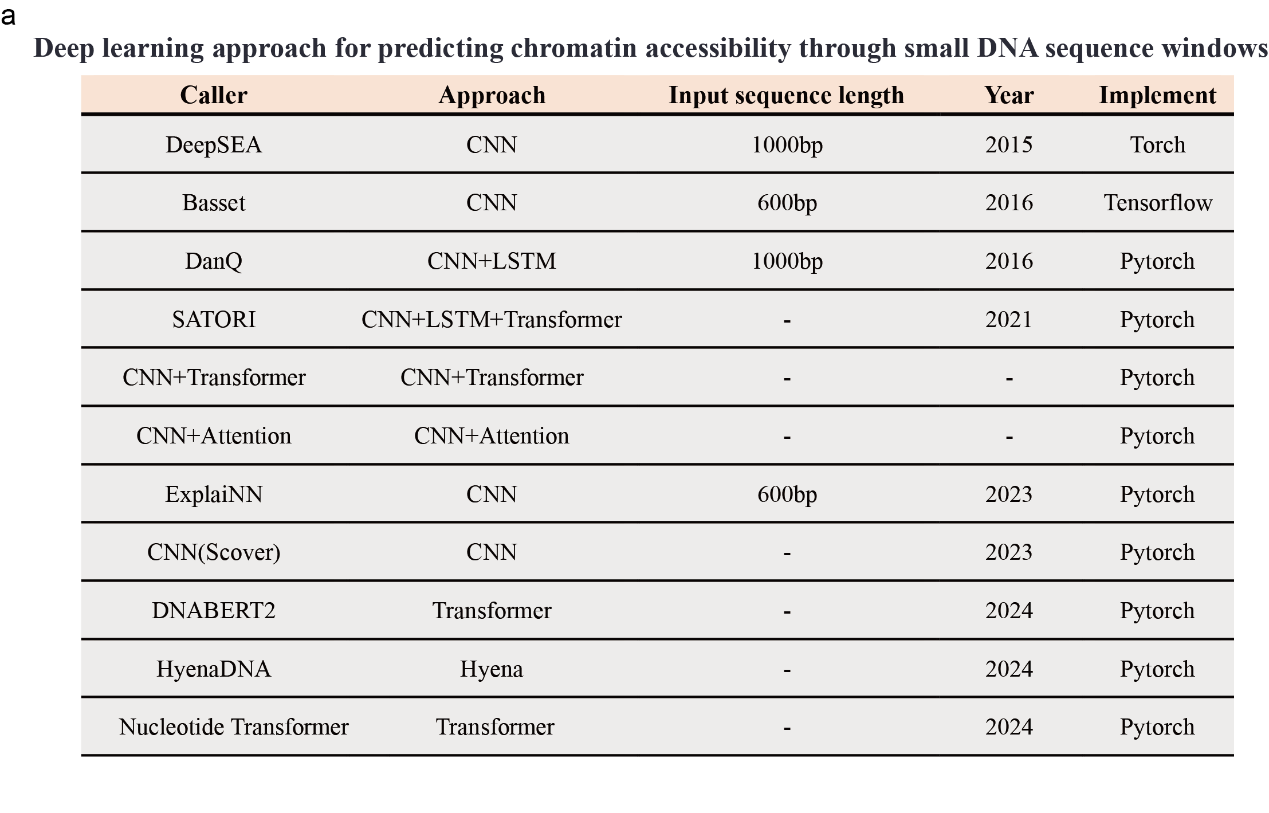

**FigS1 Sequence-function model classification. a,** The model architectures and input sequence lengths used by currently published sequence-function models, along with the year of publication and the deep learning frameworks employed.

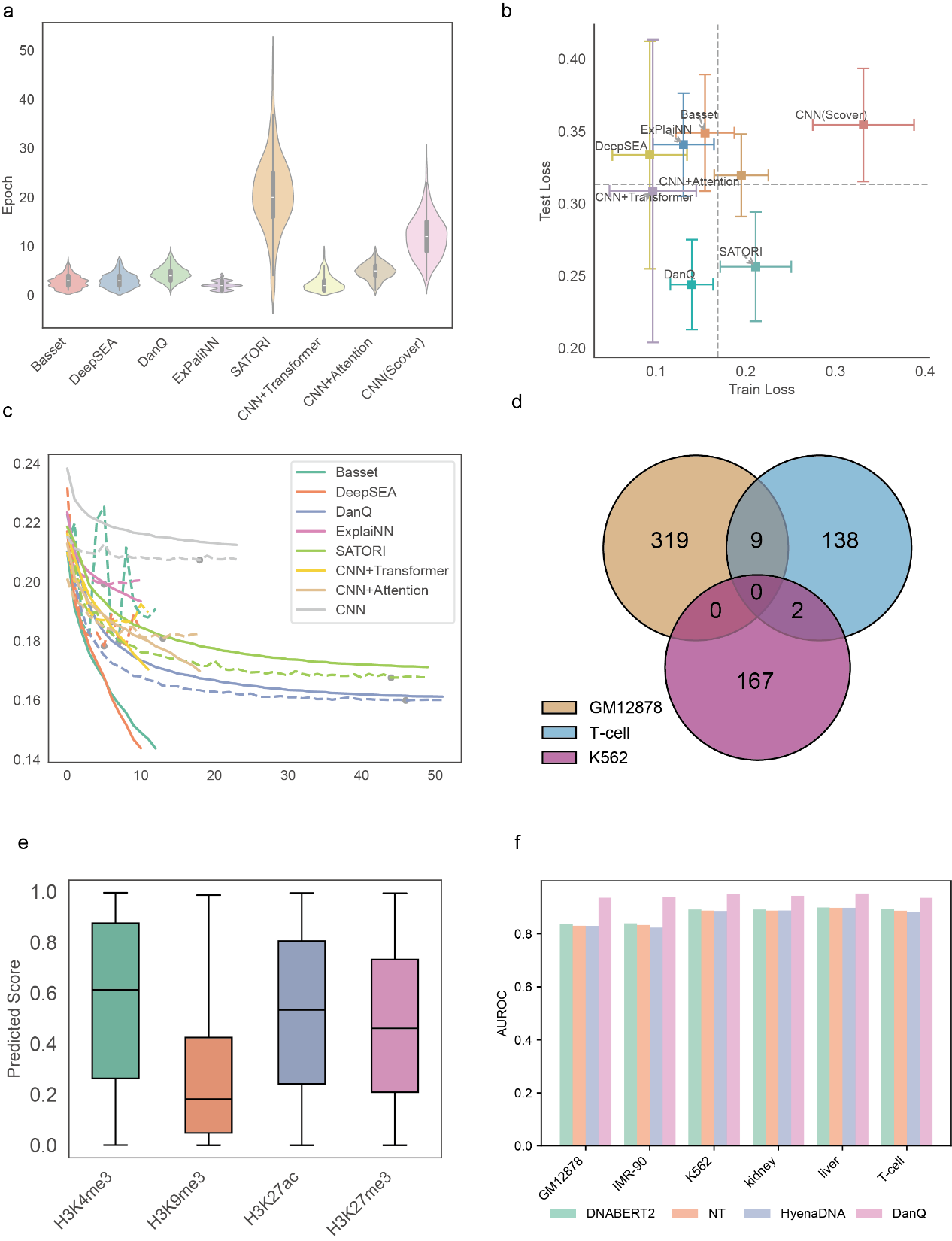

**FigS2 Performance of the algorithm in predicting binary chromatin accessibility. a,** Convergence speed of different models. **b,** Overfitting tendencies in single-task models. **c,** Overfitting tendencies in multi-task models. **d,** Overlap of regions with predicted scores less than 0.4 in different cell lines. **e,** Prediction scores of the model across regions marked by different histone modifications. **f,** Comparative performance of DNA foundation models (DNABERT2, Nucleotide Transformer, HyenaDNA) and DanQ on six cell line ATAC-seq datasets.

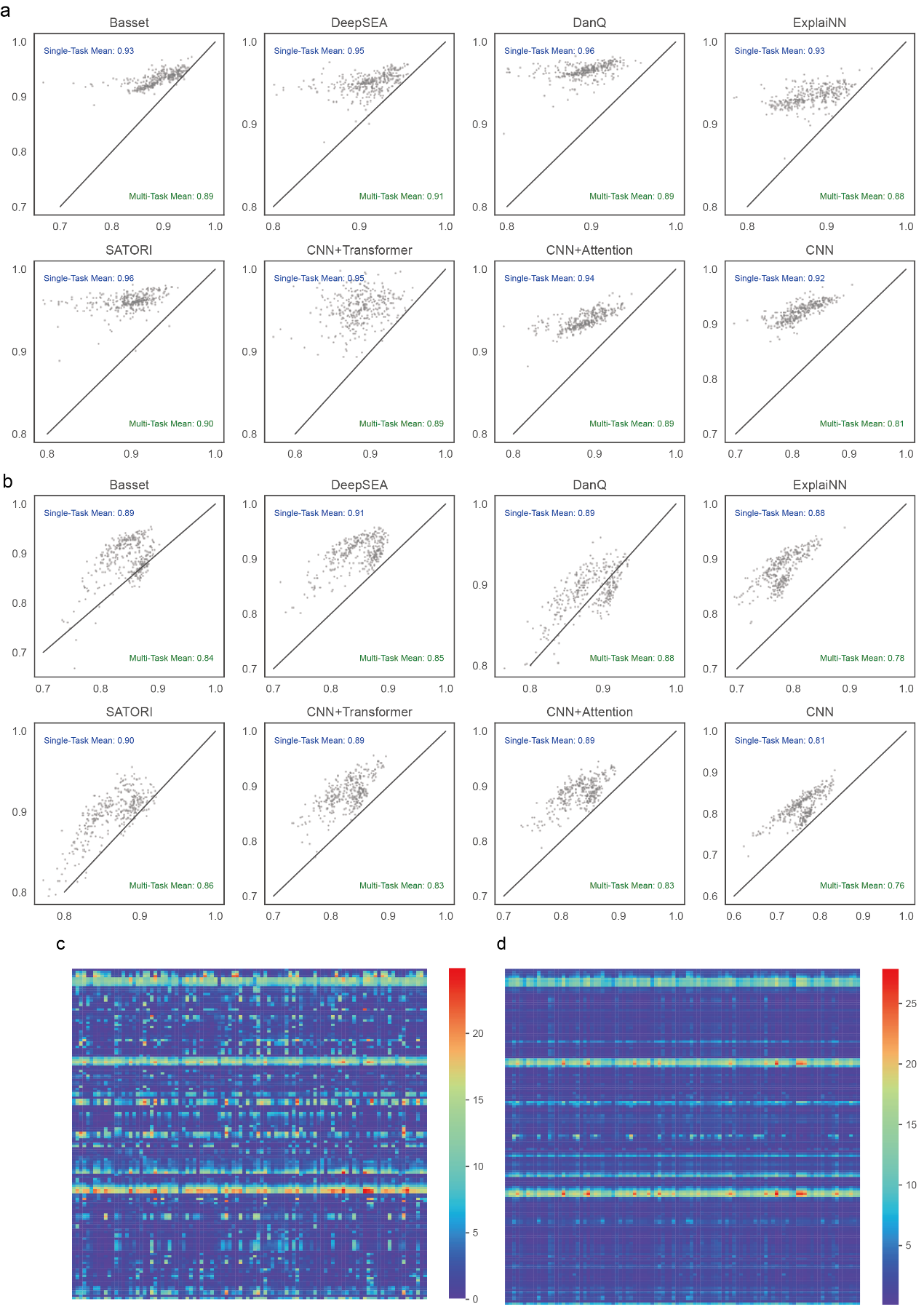

**FigS3 a,** Comparative analysis of single-task and multi-task learning models on single-task and multi-task datasets. **b**, Evaluation of single-task models and multi-task on multi-task datasets. **c,** Real Chromatin Accessibility. **d**, Chromatin accessibility predicted by the model

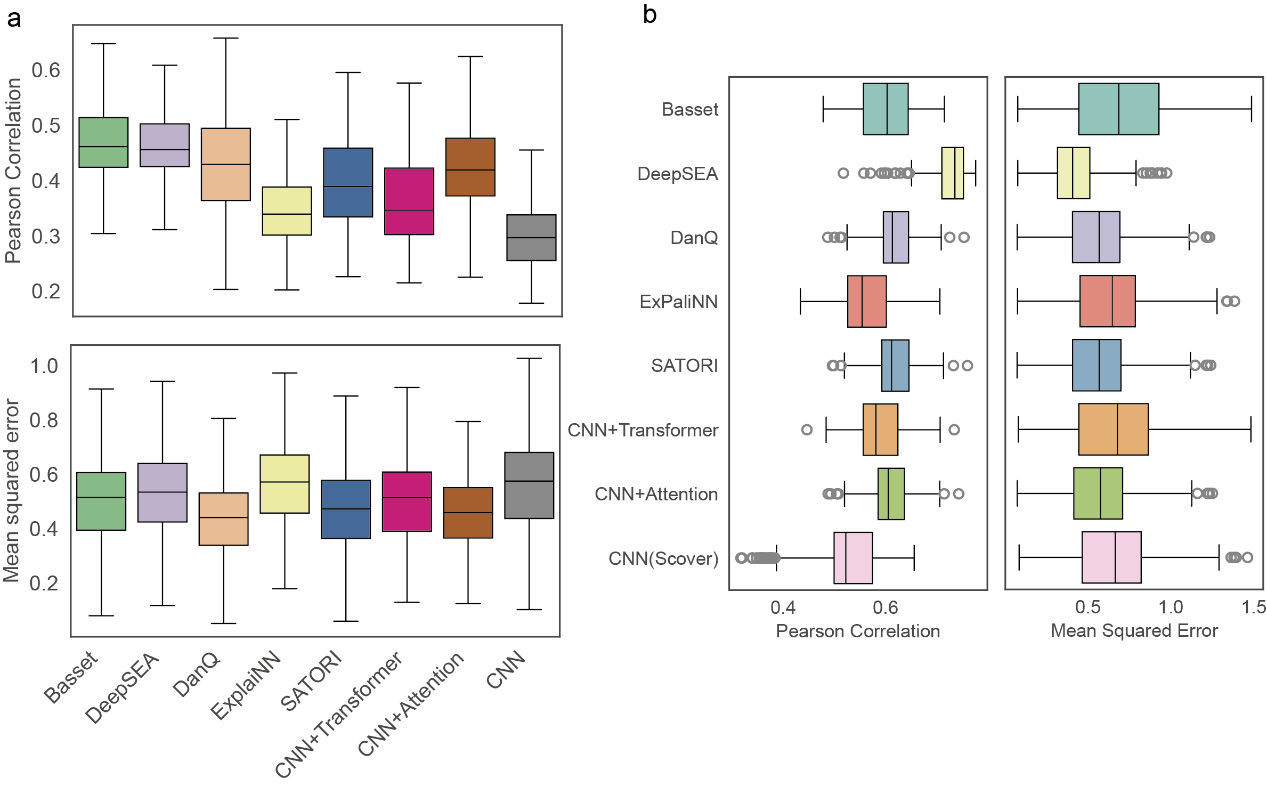

**FigS4** **a,** The Pearson correlation coefficients (PCCs) of single-task models in chromatin accessibility regression tasks. **b,** Performance of multi-task models across tissues and cell lines.

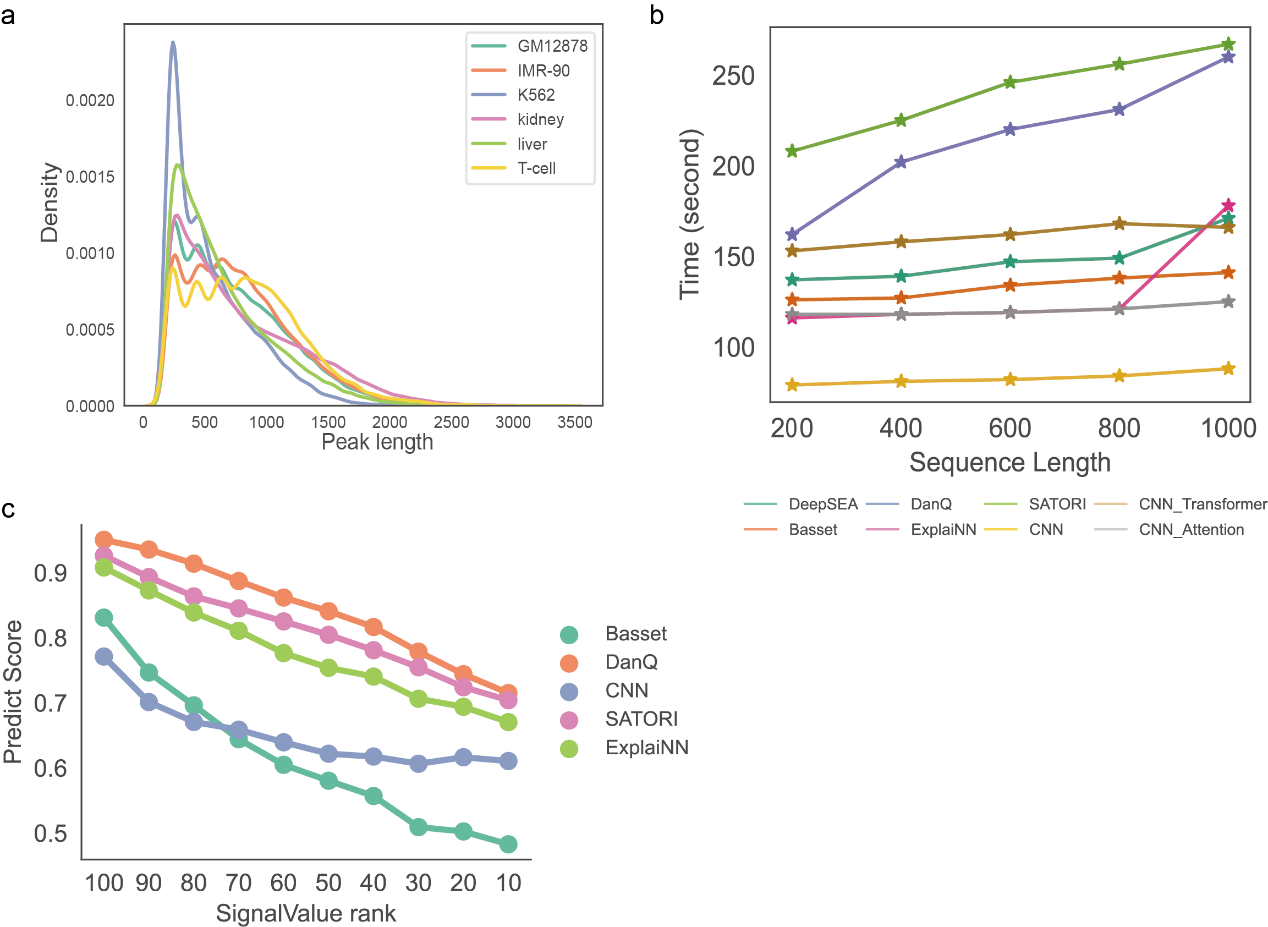

**FigS5** **a**, Distribution of peak lengths across six cell lines or tissues datasets. b, Training time variations with input sequence length, showing linear increases for LSTM-based models and stable times for non-LSTM models. c, Correlation between signal intensity and model performance.

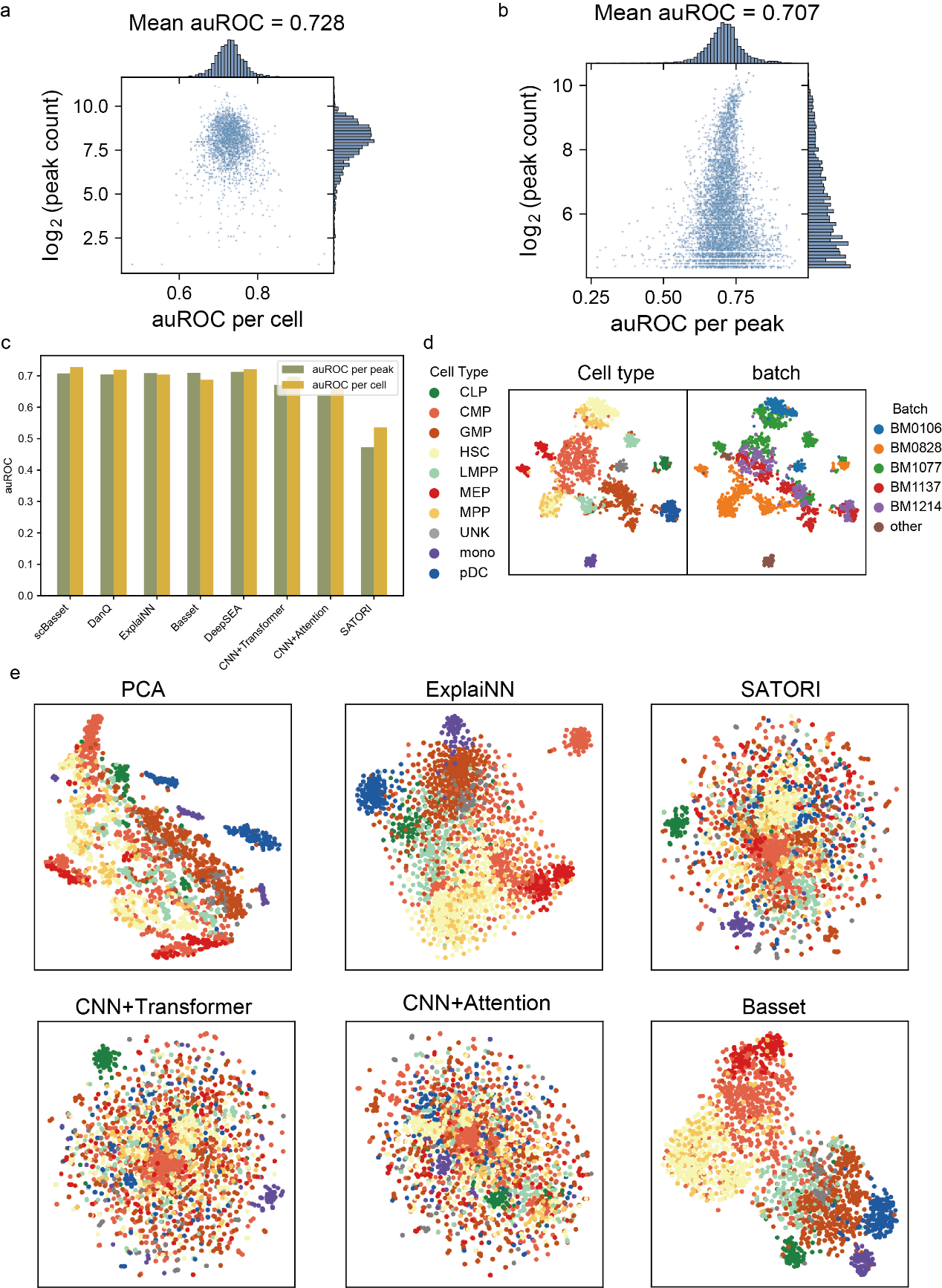

**FigS6** **a, b**, Performance of scBasset in predicting chromatin accessibility at the single-cell and peak levels. **c,** Performance of multiple algorithms in predicting chromatin accessibility at the single-cell and peak levels. **d,** t-SNE visualization of single-cell embeddings generated by scBasset. **e,** The visualization of t-SNE by other methods.

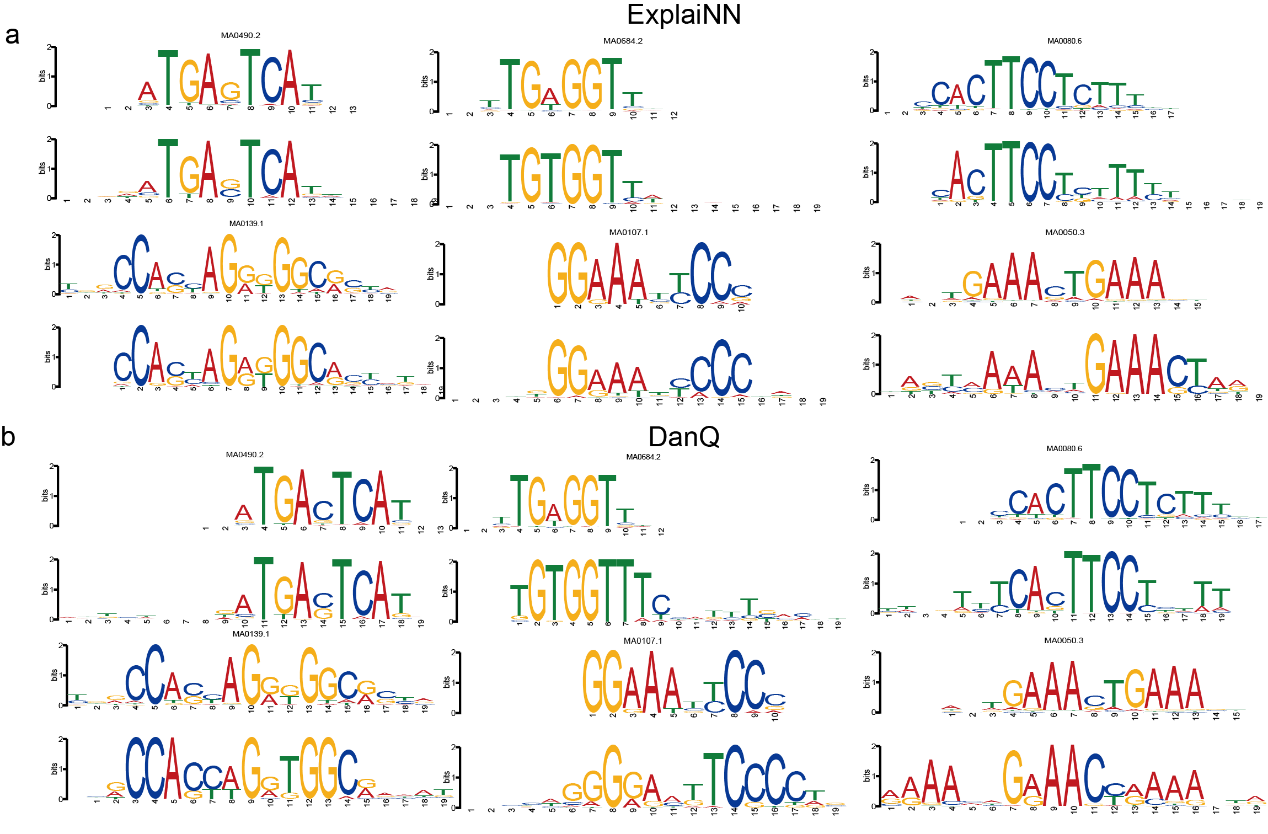

**FigS7 a,** Motifs learned from the first convolutional layer of ExplaiNN**. b,** Motifs learned from the first convolutional layer of DanQ.

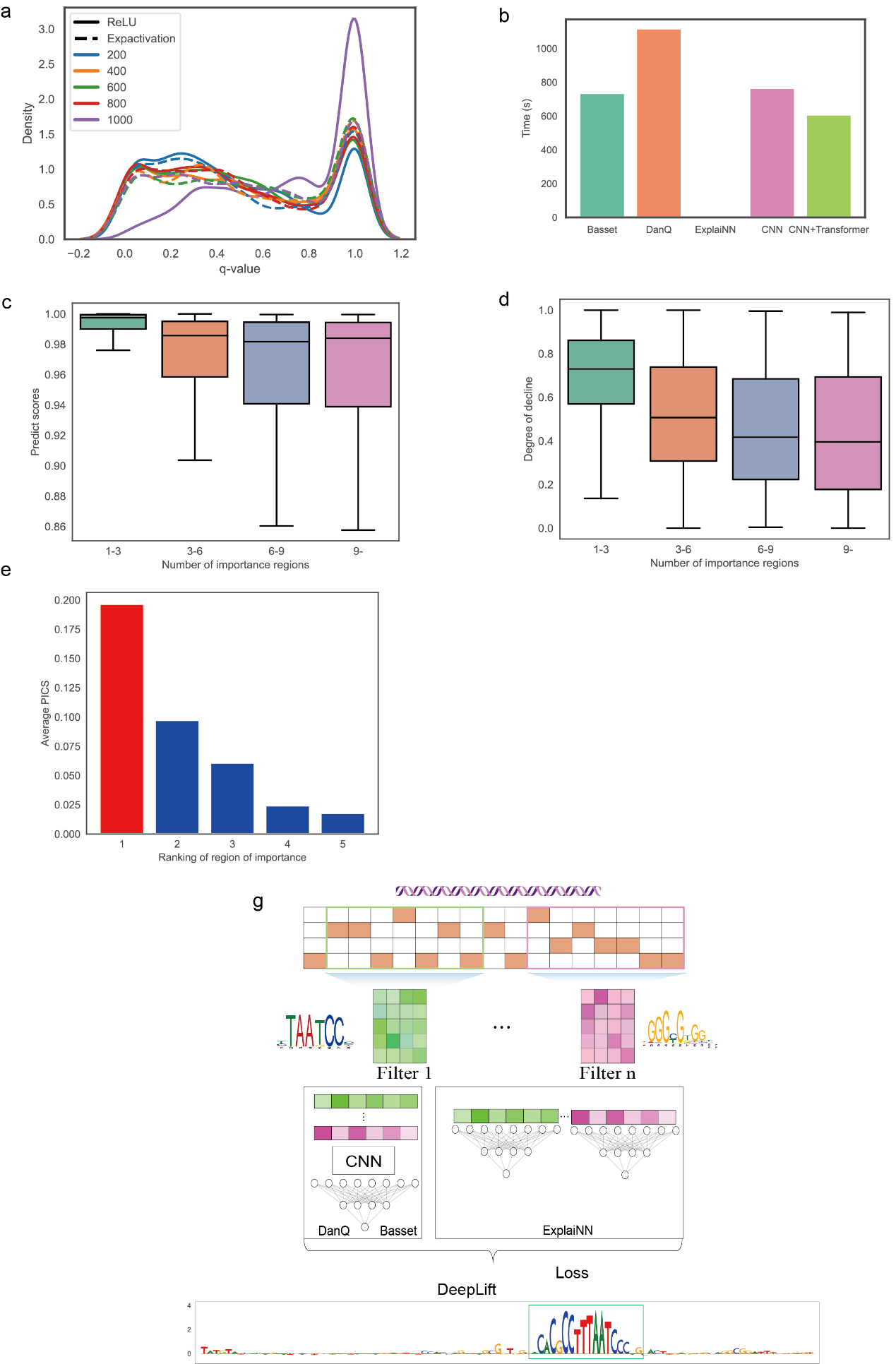

**FigS8** **a,** The motifs learned by the first convolutional layer of the DanQ model at different input sequence lengths were compared with known motifs from the JASPAR database, with q-values calculated using MEME. **b,** Time required to quantify the impact of filters in the model. **c,** Sequences contain relationships between significant regions and model predicted scores. **d,** Degree of change in the model predictions for the different attribution regions of the disturbance. **e**, Based on population fine mapping data, GWAS SNPs that may be causal have a greater probability of PICS in high significance regions than in low significance regions. **f,** The hybrid model architecture uses DanQ or Basse as the backbone and ExplaiNN as the side branches to improve the interpretability of the model.

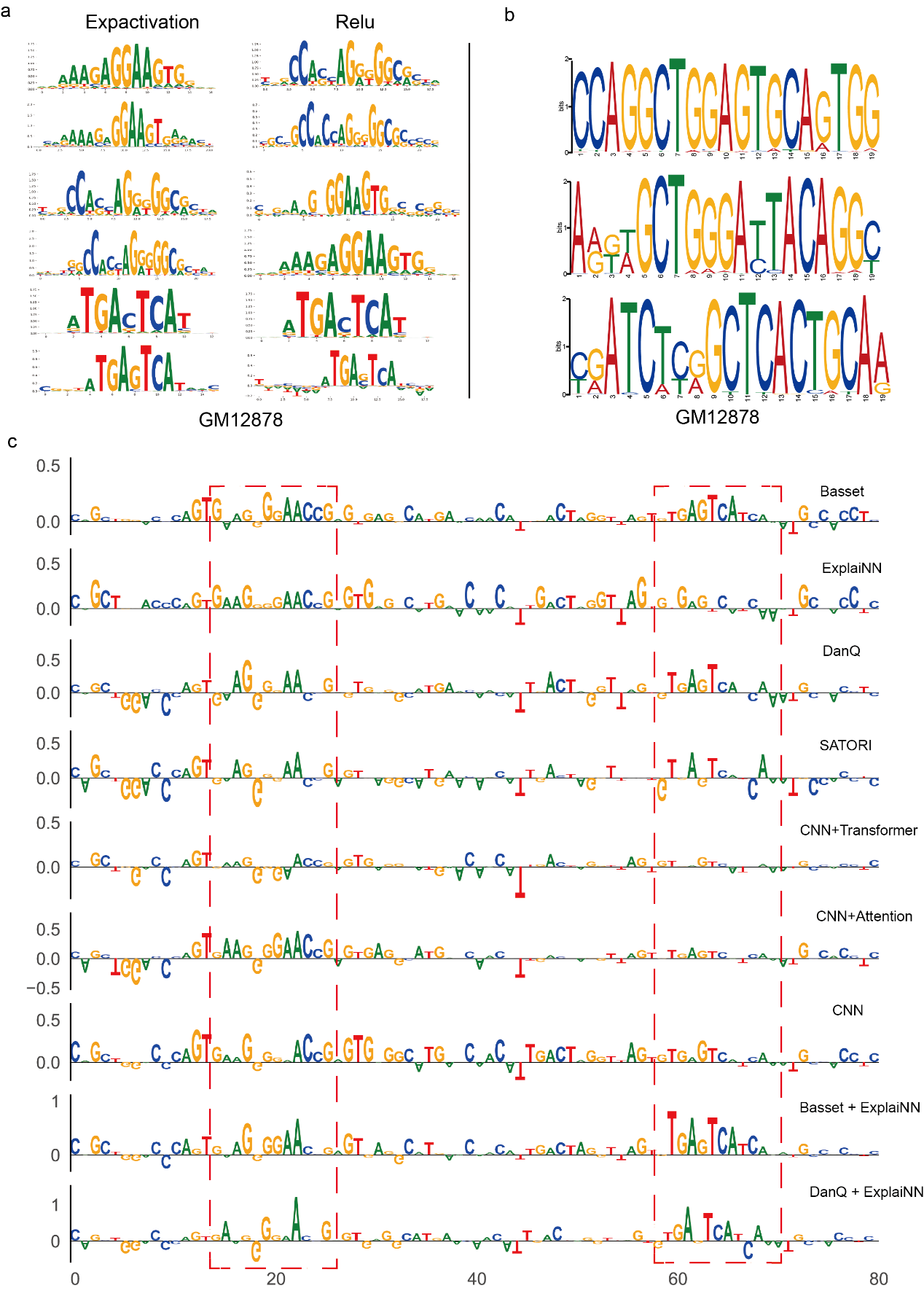

**FigS8 a,** Clustering analysis of the DeepLIFT attribution values generated by the DanQ model was performed using the TF-MoDISco tool to identify and extract relevant transcription factor binding motifs. **b**, Identification of transcription factor binding motifs from chromatin accessibility data using MEME. **c**, Attribution values of DeepLIFT across different methods.

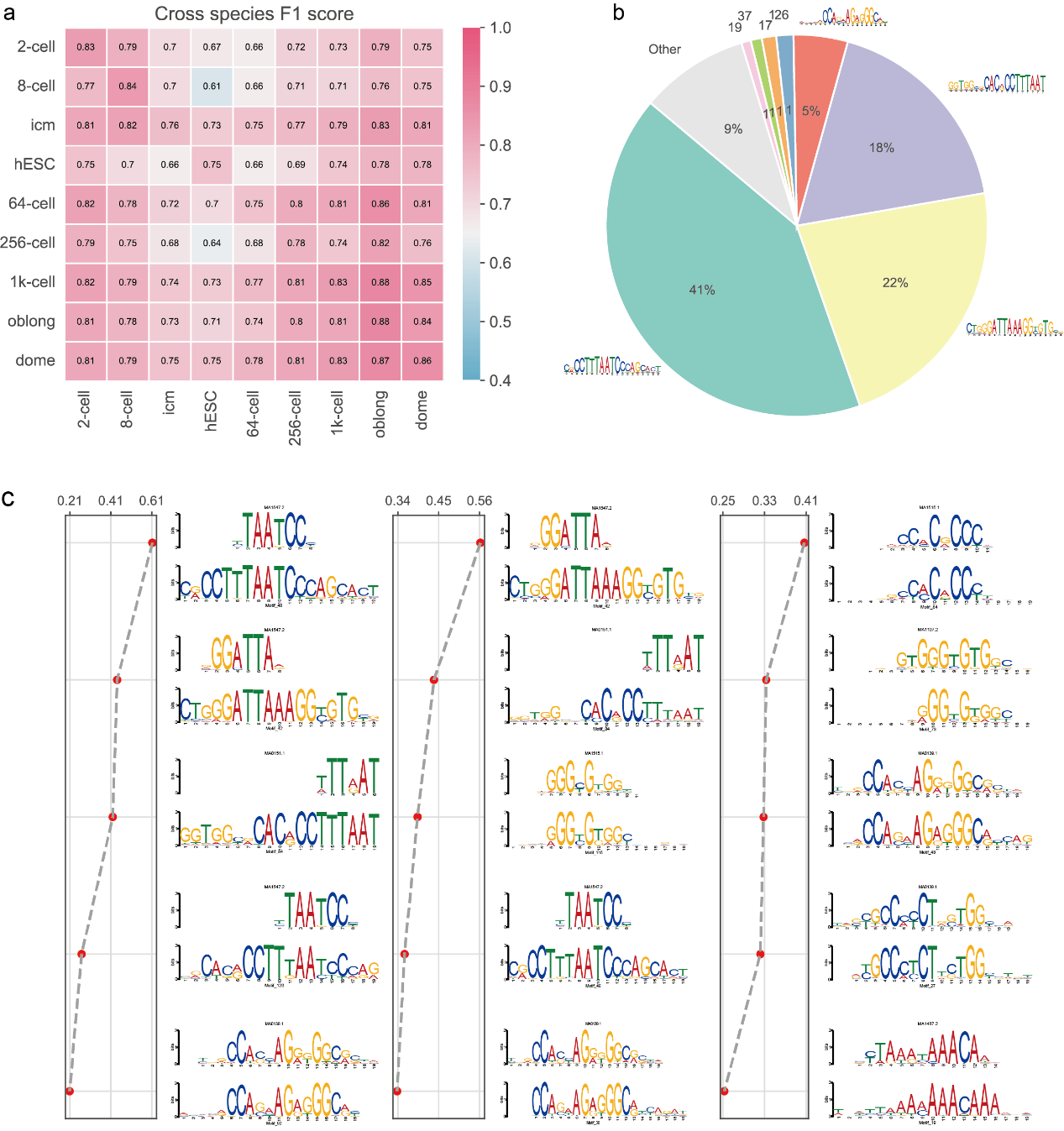

**FigS9 a,** Human and zebrafish cross-species chromatin accessibility prediction F1 score. **b,** Percentage of motifs in chromatin accessible regions found by sequence-function modelling. **c,** Sequence motifs found at different developmental stages during early mouse embryo development and the weights representing their filters (higher means more important for the model to predict as chromatin accessibility regions).

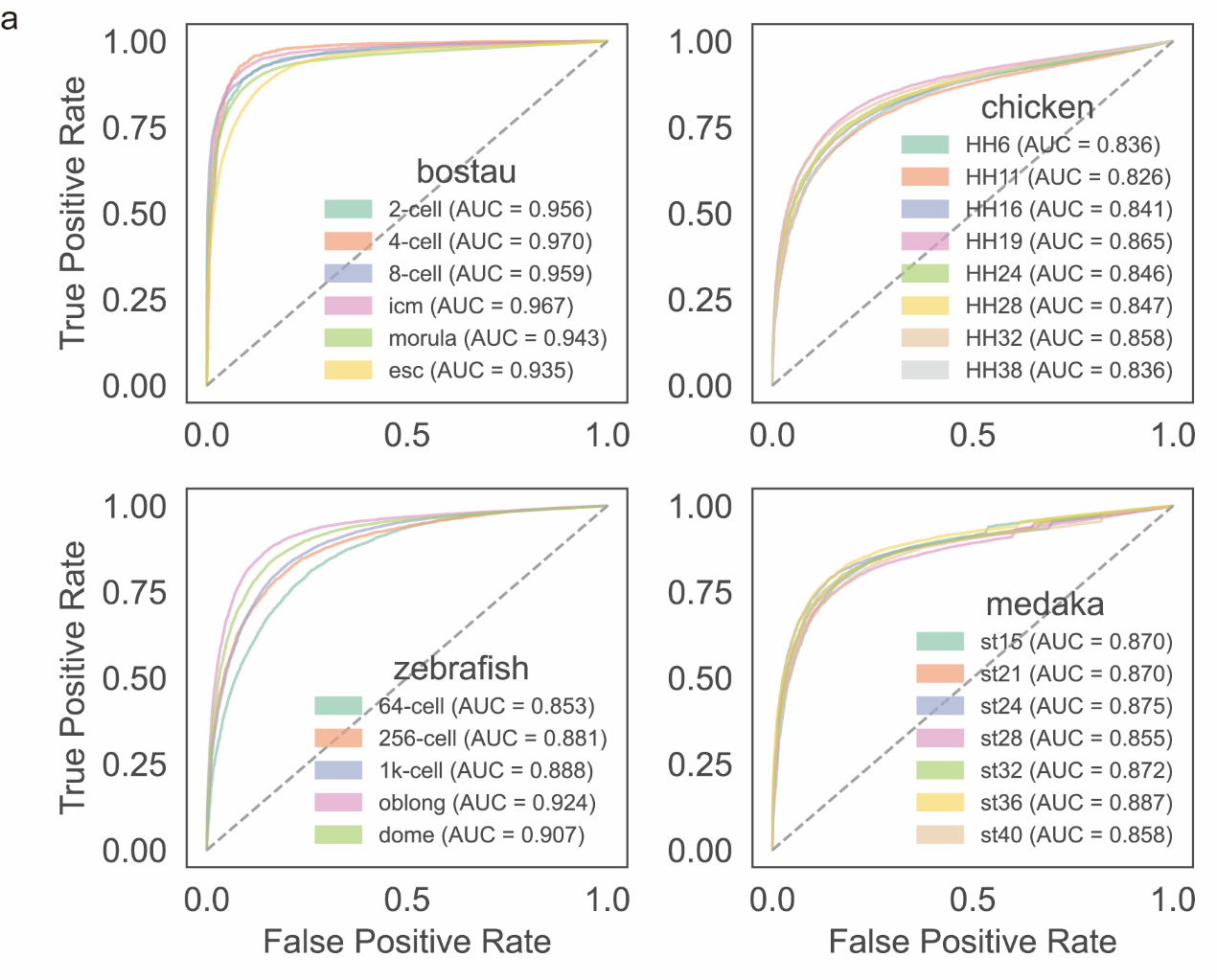

**FigS10 The AUROC for four other vertebrate species.**
